# Lipid lowering alone fails to limit atherosclerosis progression and neutrophilic inflammation in middle-aged mice

**DOI:** 10.1101/2025.11.04.686598

**Authors:** Olivia Gannon, Allison Rahtes, Jesse Bonin, Ignacia Salfate del Rio, Jessica Partridge, Sayeed Khan, Ariana Nobles, Gideon R. Covert, Amber Bahr, Ramon Bossardi Ramos, Katherine C. MacNamara, Gabrielle Fredman

**Affiliations:** Department of Molecular and Cellular Physiology, Albany Medical College, Albany, NY 12208, USA; Department of Immunology and Microbial Disease, Albany Medical College, Albany, NY 12208, USA, Albany Medical College, Albany, NY 12208, USA

**Author notes:** Correspondence should be addressed to G.F., The Department of Molecular and Cellular Physiology or K.C.M. Department of Immunology and Microbial Disease Albany Medical College, Albany Medical College, Albany, NY 12208, USA. Supported by the NIH National Heart, Lung, and Blood Institute grant HL170249 (G.F., K.C.M.). Disclosures: None Disclosed. Authors contributed equally (Olivia Gannon, Allison Rahtes).

**Keywords:** atherosclerosis, neutrophil, aging, lipid-lowering, inflammation

## Abstract

While cholesterol lowering remains the cornerstone therapy for atherosclerosis and prevention of major adverse cardiovascular events, 33–50% of individuals on lipid-lowering therapy continue to exhibit elevated inflammation. Middle age (MA) represents a critical window for disease acceleration, underscoring a need to better understand non-resolving inflammation in this time frame. We rendered young (2 months) and MA mice (10 months) hypercholesterolemic with an AAV8-PCSK9 virus and fed a western diet (WD). Following 20 weeks on a WD, mice were switched to chow for 6 weeks to mimic lipid-lowering therapy. We found that MA atherosclerotic mice had increased plaque necrosis with increased circulating and bone marrow PMN compared to young mice. Upon lipid lowering, unlike in young mice, MA mice had increased necrosis, and reduced remodeling as well as increased circulating white blood cells and bone marrow hematopoietic stem cell progenitors (HSPCs). Circulating neutrophils correlated with necrosis in MA mice whereas young mice exhibited no correlation. Lipid lowered MA atherosclerotic mice had bone marrow HSPCs and neutrophils that exhibited a more activated phenotype relative to young. Elevated neutrophil-endothelial contacts were observed in the hippocampal vasculature of MA lipid lowered mice. While lipid lowering restrains atheroprogression in young, it is inadequate in MA, failing to reduce systemic inflammation and indicating the need for complementary therapies during this time frame.

## Introduction

Cholesterol lowering remains the mainstay therapy for atherosclerosis and prevention of major adverse cardiovascular events. However, 33–50% of individuals on lipid-lowering therapy continue to exhibit elevated inflammation^1,2^. Importantly, middle age (MA) represents a critical window when disease acceleration occurs^3,4^, underscoring the need to better understand non-resolving inflammation in atherosclerosis^5^ during this critical time frame. Atherosclerosis was previously thought to be a slow progressive disease throughout one’s lifespan, but new research suggests that atherosclerotic plaques develop in an episodic manner and accelerate rapidly in some individuals between 40-50 years of age ^6,3^. This prompted us to question what cells/cellular processes are associated with middle age that may lead to rapid acceleration of plaques.

Changes in immune cell production, and enhanced myelopoiesis, is an established mechanism that underlies systemic inflammation in atherosclerosis^7^. Recent findings indicate that MA is a defining period in bone marrow health, during which hematopoietic changes are still reversible^8,9^. This suggests that examining both the bone marrow and circulating immune cells may not only reveal signs of unresolved inflammation but also provide deeper insight into mechanisms governing immune function in the context of middle age, atherosclerosis, and lipid lowering. Previous work evaluated bone marrow from old mice and observed slightly worsened atherosclerosis, and that lipid lowering conditions could not restore the function of plaque macrophages^10,11^. Given the clinical importance of the MA time frame, we posit that identifying myeloid cells linked to systemic inflammation in MA and atherosclerosis may provide insight into the cellular mechanisms driving not only plaque formation but also the broader multi-organ effects of atherosclerosis^12^.

Indeed, emerging clinical data suggests a link between atherosclerosis and cognitive decline and newer findings reveal that middle age atherosclerosis is associated with vascular dementia later in life^13–16^. Animal models of atherosclerosis exhibit signs of neuroinflammation like microgliosis and astrogliosis^17–19^. Yet gaps remain in our understanding as to whether circulating myeloid cells impact the brain during atherosclerosis and whether lipid lowering confines atherosclerosis related neuroinflammation.

Herein, we found that MA mice exhibited increased plaque necrosis and increased polymorphonuclear neutrophils (PMNs) compared to young mice after 20 weeks of western diet. Further, upon lipid lowering, MA mice had increased plaque necrosis, reduced plaque remodeling, as well as increased circulating white blood cells, and increased hematopoietic stem cell progenitors (HSPCs) in the bone marrow. Additionally, increased circulating and mature bone marrow neutrophils were directly correlated with increased plaque necrosis in MA mice on a lipid lowering diet, whereas no such relationship was observed in young mice on the same regiment. Upon lipid lowering, bone marrow from MA atherosclerotic mice had more HSCs and multipotent progenitors, and bone marrow neutrophils exhibited a more activated phenotype. Atherosclerosis in MA mice had systemic impacts, as we also observed that MA atherosclerotic mice had elevated neutrophils in the hippocampal vasculature. Overall, these results highlight that lipid lowering alone is not sufficient to limit atherosclerosis progression during middle age. The implication of our findings is that additional therapies are needed to fully attenuate disease during this critical time frame.

## Methods

### Atherosclerotic Plaque Lesion Analysis

Male C57BL/6 mice (40- or 8-week-old) were purchased from Jackson Labs (cat# 000664, Jackson Labs, Bar Harbor, ME USA) and remained in-house for the duration of the study. Mice were housed on a 12-light/ 12-dark light cycle in groups of 1-5, and provided with standard mouse chow (unless given experimental diet), water ad libitum, pinechip bedding, and environmental enrichment (nestlets and shepherd-shacks). All procedures were approved by Albany Medical Center IACUC. Mice were injected with AAV-PCSK9 (1x10^12^ MOI) and immediately placed on Western Diet (Teklad Diets, cat# TD.88137, Inotiv, West Lafayette, Indiana, USA). Mice were randomly assigned to receive 20-weeks of WD diet feeding (i.e. baseline) or 20 weeks of WD followed by a chow diet switch for an additional 6 weeks (i.e. lipid lowering). Plasma Cholesterol was measured using the cholesterol E kit from Fujifilm (cat# 999-02601, Fujifilm, Lexington, MA, USA) following the manufacturer’s instructions. Briefly, blood was collected via retro-orbital collection using a heparinized capillary under sedation just prior to euthanasia.

Lesion and necrotic area were measured as previously described^20^. In brief, hearts were perfused with PBS and aortic roots were frozen in OCT at the time of harvest. Aortic roots were sectioned starting at the base of the plaque in 10 µM increments, with two sections per glass slide for a total of approximately 30 sections. Sections were stained with Hematoxylin & Eosin (H&E) and analysis was performed using an average of 6 slides for a total span of ∼100 µm to capture the entire plaque. Lesion area was defined as the region from the internal elastic lamina to the lumen.

### Atherosclerotic Plaque Immunofluorescence

Frozen aortic root sections were fixed with either acetone (Mac2, αSMA) or methanol (H3Cit) prior to blocking (1% BSA, 10% goat serum, 1 drop M.O.M. (cat# MKB-2213-1, Vector Labs, Newark, CA, USA) in DPBS). In addition, for Mac2 and αSMA stains, sections were permeabilized in a solution containing 0.01%-0.05% Triton X-100 (cat# 9036-19-5, Sigma Aldrich Inc, St. Louis, MO, USA) prior to blocking. After brief washes, samples were stained with either anti-Mac2 (cat# CL8942AP, Cedarlane Labs, Burlington, ON, USA) at 1:10000, anti-Histone H3 (citrulline R2 + R8 + R17) (cat#AB281584, clone RM1001, Abcam, Waltham, MA, USA) at 1:500, or anti-smooth muscle actin (cat#. 23081-1-AP, Proteintech, Rosemont, IL, USA) at 1:1000 in 1%BSA. Secondary antibodies used were Alexa Fluor 594 anti-rat (cat# A-11007, Thermo Fisher Scientific, Carlsbad, CA, USA) (Mac2) and Alexa Fluor 594 anti-rabbit (cat# A-11037, Thermo Fisher Scientific, Carlsbad, CA, USA) (H3Cit and αSMA). Secondary was applied at 1:200 (Mac2, H3Cit) for 2 hours at 4°C or 1:1000 (αSMA) for 1 hour at room temperature. Nuclei were stained with Hoescht (cat# H3750, Thermo Fisher Scientific, Carlsbad, CA, USA). Sections were rinsed 3X in DPBS after incubation with primary, secondary, and Hoescht and stored in PBS at 4°C until imaging. All sections were imaged within a day or two of initial staining and were imaged using a Lecia Thunder Imaging system and LAS X software. Lesional Mac2+ cells were determined using HALO AI software version 8.1 and the Area Quantification FL protocol (HALO, Indica Labs, Albuquerque, NM, USA). H3Cit MFI and total lesion area were determined using ImageJ software to determine % area H3Cit. ImageJ was also used for the analysis of cap αSMA MFI. All values were compared for statistical significance using Graphpad Prism v10.1 software and the two-way ANOVA with Tukey’s multiple comparison test.

### Complete Blood Counts

Blood was collected retro-orbitally using ammonium heparinized microhematocrit capillary tubes (cat# 51608, Globe Scientific, Mahwah, NJ, USA) into EDTA-coated vials (cat# 365974, Globe Scientific, Mahwah, NJ, USA) and were analyzed for complete blood counts (CBC) on a Heska Element HT5 (Heska, Loveland, CO, USA).

### Cell Preparation and Flow Cytometry

Hind limbs and spleens were collected at the time of euthanasia and processed as previously described^21^. Red blood cells were lysed, and cells were counted using an Invitrogen Countess 3 (ThermoFisher, Waltham, MA, USA). Single cell suspensions were plated, and Fc receptors were blocked (anti-CD16/32 (clone 2.4G2) antibody (cat # 553141, BD Biosciences, Franklin Lakes, NJ, USA). Cells were then stained with antibody cocktails (**Table 1**), followed by secondary staining when applicable, and fixed in 1% paraformaldehyde solution. Data were acquired on a Cytek Northern Lights flow cytometer and SpectroFlo software (Cytek Biosciences, Fremont, CA, USA) and analyzed using FlowJo software version 10.7.1 (TreeStar, Ashland, OR, USA). For assays examining responses to PMA stimulation, bone marrow cells were isolated from the hind limb as described above and 3x10^6^ cells were incubated with either vehicle or 160nM phorbol 12-myristate 13-acetate (P1585, Sigma-Aldrich, Burlington, MA, USA), for 2 hours at 37°C, as previously done ^22^, and stained and analyzed by flow cytometry.

**Table 1:** List of antibodies used for Flow cytometry analyses.

| Marker | Clone | Fluorophore | Supplier | Catalog Number |
| --- | --- | --- | --- | --- |
| CD11b | M1/70 | APC-Cy7 | BioLegend | 101226 |
| CD3 | 17A2 | APC-Cy7 | BioLegend | 100221 |
| CD45 | RA3-6B2 | APC-Cy7 | BioLegend | 103224 |
| Ter-199 | TER-119 | APC-Cy7 | BioLegend | 116223 |
| Ly-6G/Ly-6C (Gr-1) | RB6-8C5 | APC-Cy7 | BioLegend | 108242 |
| Streptavidin | - | BV785 | BioLegend | 405249 |
| CD135 | AF210 | PE | BioLegend | 135306 |
| CD150 | TC15-12F12.2 | BV711 | BioLegend | 115941 |
| CD48 | HM48-1 | Pacific Blue | BioLegend | 103418 |
| cKit | 2B8 | APC | BioLegend | 105812 |
| Sca-1 | D7 | PE-Cy7 | BioLegend | 108114 |
| Ly6G | 1A8 | BV605 | BioLegend | 127639 |
| Ly6G | 1A8 | SparkNIR 685 | BioLegend | 127665 |
| Ly6C | HK1.4 | Pacific Blie | BioLegend | 128013 |
| CD63 | NVG-2 | AlexFluor 700 | BioLegend | 143924 |
| SiclecF | S17007L | Biotin | BioLegend | 429404 |
| 7-AAD | - | PerCP Cy5.5 | BioLegend | 155532 |
| **Table 1: List of antibodies used for Flow cytometry analyses** |  |  |  |  |

### Colony Forming Unit (CFU) Assays

Bone marrow and spleen cells were isolated and single cell suspensions were prepared as described above. Cells were plated at a density of 20,000 cells/well (bone marrow) or 100,000 cells/well (spleen) in 35mm dishes in 1 mL of Methylcellulose-based medium (cat# GF M3434, Stemcell Technologies, Cambridge, MA, USA). Cells were maintained in a humidified incubator at 37°, 5% CO2 for 6-8 days and then colonies were identified as CFU-G (granulocyte), CFU-M (macrophage), CFU-GM (granulocyte macrophage) or CFU-GEMM (granulocyte erythroid macrophage megakaryocyte) and counted.

### Bone Marrow Immunohistochemistry

Humerus and calvaria bones were dissected, cleaned of excess tissue, and drop-fixed in 10% neutral buffered formalin (cat# HT501320-9.5L, Sigma-Aldrich, Louis, MO, USA) for 48 hours at room temperature followed by decalcification in 14% EDTA solution (cat# 118430050, Thermo Scientific, Waltham, MA, USA) for 14 days with agitation. Bones were then embedded in paraffin, sectioned (5µM), and stained with primary antibodies against Ly6G (cat# 551459, BD Pharmingen, BD Biosciences, Franklin Lakes, NJ, USA) followed by secondary antibodies (cat# BA-9401, Vector Labs, Newark, CA, USA). At least 2 sections were imaged per mouse at 10x magnification. ROIs were drawn by a blinded observer in ImageJ, the number of Ly6G+ cells were counted, normalized to the ROI size, and averaged.

### RNA-Seq

Total bone marrow was collected from hind limbs and red blood cells were lysed followed by resuspension in 1mL TRIZol. Bulk RNA-seq datasets are available in NCBI’s GEO under accession numbers GSE308089. RNA-seq libraries were prepared using Illumina TruSeq protocols and sequenced on the Illumina NextSeq 500. Reads were aligned to mm10 (mouse) reference genomes with Rsubread (v2.10.5), and gene counts were obtained using featureCounts. Genes with CPM > 0.5 were retained and annotated to gene symbols using NCBI. Count matrices were normalized with the TMM method in edgeR, log2-CPM transformed, and analyzed with the limma-voom pipeline. To infer transcription factor (TF) activity from bulk RNA-seq data, we implemented the DoRothEA framework. Enrichment scores (NES) > 2 were considered indicative of TF activation. This analysis enabled the identification of key regulators underlying differential transcriptional states.

### ATAC-Seq

Bone marrow cells were isolated from the hind limb, red blood cells were lysed and enumerated. 1*10^5^ cells were flash frozen in 10% DMSO. Experiments were performed at the Cornell BRC Epigenomics Facility using the Omni ATAC-seq protocol with 100k cells, followed by 12 cycles of PCR amplification. Libraries were sequenced on the Element AVITI platform (>10M paired-end reads/sample). Reads were filtered to remove mitochondrial (chrM) and ENCODE blacklisted regions. Peaks were called with MACS3 (q < 0.1), and FRiP scores were computed for quality control. Coverage tracks were generated with bamCoverage (CPM normalization). Consensus peak sets were constructed with bedtools merge across replicates, and peak counts were quantified with bedtools multicov.

### Immunofluorescence of brain Ly6G-Lectin staining

Brains were collected at the time of harvest (post perfusion with PBS) and fixed via submersion in 4% paraformaldehyde overnight at 4°C. Following fixation, brains were submerged in 30% sucrose in DPBS for 48 hours at 4°C prior to embedding in optimal cutting temperature compound and storage at -80°C. Brains were sectioned into 40 μm cross sections using a Leica CM1950 UV Cryostat (Leica, Teaneck, NJ, USA) and stored in cryopreserve compound (500 mL 1XPBS, 10g Polyvinyl-pyrrolidone mol wt. 40,000, 300 mL Ethylene Glycol, 300 g sucrose, 0.1g Sodium Azide) at -20°C until the time of staining. Prior to staining, cryopreserve compound was removed with 1XPBS in a 15mm diameter Netwell insert (cat# 3477, Corning, Glendale, AZ, USA). Sections were then permeabilized (0.5% triton in 1XPBS), and blocked (0.5% triton and 4% donkey serum in 1XPBS) at room temperature prior to incubation with primary antibody solution (Ly-6G Monoclonal Antibody, clone 1A8-Ly6g (cat# 16-9668-82 ThermoFisher, Waltham, MA, USA) at 1:250 in 1XPBS for 24 hours at 30C. After 24 hours, sections were incubated with a secondary antibody solution containing Alexa Fluor 594 anti-rat antibody (cat# A-11007, Thermo Fisher Scientific, Waltham, MA, USA) at 1:300 and a fluorescein-conjugated Lectin antibody (cat# FL-1171-1, Vector Laboratories, Newark, CA, USA) at 1:100 for 2 hours at 30C.Stained brain sections were visualized on a Leica Thunder Imager using LAS X software. Lectin and Ly6G colocalization areas were analyzed by creating a Lectin area ROI in desired regions of the brain section using ImageJ software version 1.54f (https://imagej.net/ij/index.html). After ROI was created and saved for different regions of the brain, the ImageJ Colocalization plugin (https://imagej.net/ij/plugins/colocalization.html) was used to find the pixels matching both Ly6G and Lectin in the images analyzed. A measurement using the Lectin ROI and the image obtained from the Ly6G-Lectin colocalization 8-bit point resulted in the Area% of Ly6G expression in Lectin.

### LC-MS/MS Metabololipidomics

*Solid Phase Extraction for LC-MS/MS*: All samples underwent solid phase extraction prior to LC-MS/MS analysis using SPE C18 columns from Biotage (cat# 220-0010-B, Biotage, Charlotte, NC, USA) and a manual extraction manifold. In brief, columns were pre-conditioned with 1 column volume of methanol followed by 1 column volume of ddH2O. Before extraction, 100pg of deuterium-labeled standards (Cayman Chemical, Ann Arbor, MI, USA) (**Table 2**) were added to each sample to assess sample recovery. Methanol-suspended sample supernatants were diluted 10:1 with pH 3.5 ddH2O before loading onto equilibrated SPE columns. Columns were then washed with 1 column volume of neutral ddH2O followed by an equivalent volume of hexane. Samples were eluted in 2 fractions: 750 µL of Methyl formate, followed by 750µL Methanol, and fractions were pooled before drying. Samples were dried under a nitrogen stream using a Turbovap LV system (cat# 415000, Biotage, Charlotte, NC, USA) and pellets were re-suspended in 50 µL of 50:50 water:methanol.

**Table 2:** List of internal standards used for LC-MS/MS.

| Internal Standard Name: | Supplier & Catalog #: |
| --- | --- |
| PGE2-d4 | Cayman #314010 |
| LTB4-d4 | Cayman #320110 |
| LXA4-d5 | Cayman #10007737 |
| 5-HETE-d8 | Cayman #334230 |
| 12-HETE-d8 | Cayman #334570 |
| 15-HETE-d8 | Cayman #334720 |
| RvE1-d4 | Cayman #10009854 |
| RvD2-d5 | Cayman #11184 |
| RvD3-d5 | Cayman #19512 |
| **Table 2: List of internal standards used for LC-MS/MS** |  |

*LC-MS/MS*: Samples were analyzed using an AB SciEx QTrap 7500 spectrometer equipped with an Exion LC system, which was fitted with a Kinetex 2.6 µm Polar C18 LC column (100 X 3 mm 2.6 µm 100 Å) from Phenomenex (cat# 00D-4759-Y0, Phenomenex, Torrance, CA, USA). Samples were separated using a gradient of water:metOH:formic acid ranging from 55:45:0.001 (v:v:v) to 2:98:0.001 at a flow rate of 0.5 mL/min. Unless otherwise noted, an injection volume of 30 µL was used for all samples. Targeted lipid mediators (**Table 3)** were monitored and quantified using an MRM-IDA-EPI scanning mode with signature Q1 and Q3 ion pairs for each molecule.

**Table 3:** List of targeted compounds for LC-MS/MS analysis and their respective internal standard.

| **Target Compounds** | **Internal Standard** |
| --- | --- |
| PGE2 | PGE2-d4 |
| PGD2 | PGE2-d4 |
| PGF2a | PGE2-d4 |
| TXB2 | PGE2-d4 |
| LTB4 | LTB4-d4 |
| 20-OH-LTB4 | LTB4-d4 |
| LXA4 | LXA4-d5 |
| 15(R)-LXA4 | LXA4-d5 |
| LXB4 | LXA4-d5 |
| 12-HHTrE | PGE2-d4 |
| 5-HETE | 5-HETE-d8 |
| 11-HETE | 12-HETE-d8 |
| 12-HETE | 12-HETE-d8 |
| 15-HETE | 15-HETE-d8 |
| 20-HETE | 15-HETE-d8 |
| RvE1 | RvE1-d4 |
| RvE2 | LXA4-d5 |
| RvE4 | LXA4-d5 |
| LXA5 | RvE1-d4 |
| 5-HEPE | 5-HETE-d8 |
| 11-HEPE | 12-HETE-d8 |
| 12-HEPE | 12-HETE-d8 |
| 15-HEPE | 15-HETE-d8 |
| 18-HEPE | 15-HETE-d8 |
| RvD1 | LXA4-d5 |
| 17(R)-RvD1 | LXA4-d5 |
| RvD2 | RvD2-d5 |
| RvD3 | RvD3-d5 |
| 17(R)-RvD3 | RvD3-d5 |
| RvD4 | LXA4-d5 |
| RvD5 | LTB4-d4 |
| PD1 | LTB4-d4 |
| 10(S),17(S)DiHDHA | LTB4-d4 |
| Maresin 1 | LTB4-d4 |
| Maresin 2 | LTB4-d4 |
| 4-HDHA | 5-HETE-d8 |
| 7-HDHA | 5-HETE-d8 |
| 13-HDHA | 12-HETE-d8 |
| 14-HDHA | 12-HETE-d8 |
| 17-HDHA | 15-HETE-d8 |

Spray voltage was 1700, collision energy was -22, source temperature was 350° C, and threshold was set to 1000 cps. Identification of lipid mediators was based on a combination of LC retention time, signal:noise of >10, and concentration above our LLOQ. If a sample did not meet these criteria, it was labeled as “not detected” (ND). Where applicable, for statistical analyses values of ND were analyzed as a value of 0. Quantification of sample lipid mediators was determined by the MRM peak area ratio and a standard calibration curve corresponding to that particular compound ranging from 0.005 to 100 pg/µL. The linear calibration curve for each compound had r^2^ values ranging from 0.98 to 0.99. In the case for which an exact corresponding internal standard was unavailable for a particular compound, one of similar physical properties was used (**Table 3**). Detection limit for most compounds ranged from 0.1 to 0.9 pg/µL.

### Statistical Analysis

Analysis and graph generation including correlation matrices and VIP plots were performed using Graphpad Prism (v10.1). When evaluating both the effects of middle-age and atherosclerosis, 2-way ANOVA were performed with Tukey’s multiple comparison test. When solely evaluating the effect of either age or treatment, we performed t-tests.

## Results

### Increased lesion necrosis in response to hypercholesterolemia in middle age

To closely mimic the development of atherosclerosis observed in in middle-aged (MA) humans^3,4^, we injected MA (10 mos.) and young control (2 mos.) C57BL/6 mice with the gain of function AAV8-PCSK9 (or AAV8-gof-POCSK9) and placed them on a western diet for 20 weeks to induce atherosclerosis (**Figure 1A**). To directly compare development of atherosclerosis in young and MA mice, cohorts were sacrificed to establish “baseline” atherosclerosis at 20 weeks. To model lipid-lowering therapy, similar cohorts were fed WD for 20 weeks then had their diet switched to chow for 6 weeks. Middle-aged (MA) mice had significantly higher body weight compared with young mice, but no differences in body weight were observed between baseline and lipid lowered (LL) groups for either age group (**Figure S1A**). Fasting blood glucose levels were also measured and MA mice had elevated fasting blood glucose compared with young baseline atherosclerotic mice (**Figure S1B**). Total cholesterol levels were not different between young and MA baseline and cholesterol levels were significantly lowered upon diet switch to chow in both young and MA mice (**Figure S1C**). Therefore, age did not impact cholesterol values at baseline and advanced age did not impair lipid lowering by chow switch.

**Figure 1.**
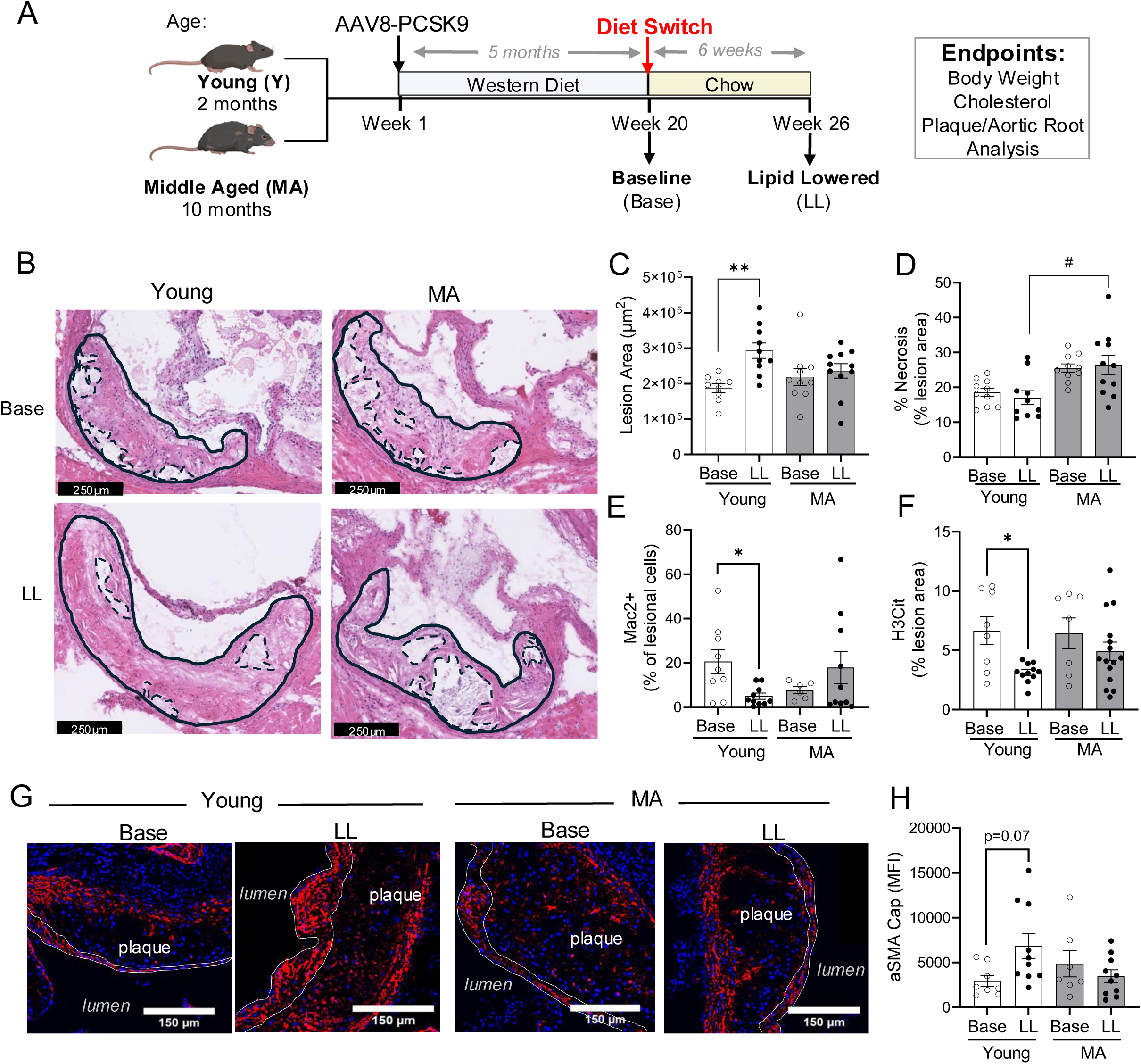
Lipid lowering in middle-aged atherosclerotic mice is associated with increased necrosis and decreased plaque remodeling compared with young controls. **(A)** The experimental design is displayed, image created with Biorender.com. **(B)** Lesion analysis of aortic root for both young and middle-aged (MA) mice after 20 weeks of western diet at either baseline (Base) or after an additional 6 weeks of lipid lowering (LL). Lesion area is indicated as a solid line, while necrotic area is outlined in dashed lines. **(C,D)** Quantification of lesion area **(C)** or percent necrosis **(D)** in Baseline (Base) and Lipid lowered (LL) young or middle-aged (MA) aortic root plaques. **(E)** Quantification of percentage lesional Mac2+ cells quantified using the HALO Area Quantification FL protocol. **(F)** Quantification of plaque percent H3Cit MFI (as a percentage of total lesion area) using ImageJ. **(G)** Representative images of alpha smooth muscle actin (αSMA, red) in the aortic roots of young or middle-aged (MA) mice at baseline or lipid lowering.**(H)** Quantification of αSMA MFI present in the lesion cap of aortic roots of young or MA mice at either baseline or lipid lowering. Results are mean ± SEM. For all analyses, each dot represents an individual mouse and comparisons were performed using two-way ANOVA with Tukey’s multiple comparison. *p<0.05, **p<0.01, effect of lipid-lowering, #p<0.05 effect of MA.

Plaque histological analysis was performed and representative images from young LL mice revealed more eosin and acellular areas, relative to baseline, which is suggestive of plaque remodeling (**Figure 1B**). Quantification revealed that lesion area was increased in young LL mice compared with young baseline controls whereas no change in plaque size was noted in MA mice upon LL (**Figure 1C**). This increase in lesion area in young mice upon diet switch is consistent with literature^23^ which suggests that lipid lowering in young mice can lead to dynamic plaque remodeling. While plaque necrosis was not significantly changed by diet switch in either age group, MA mice had increased necrosis compared with young mice after lipid lowering (**Figure 1D**). These data suggest that atherosclerosis progressed in a maladaptive manner in MA mice, despite lipid lowering. Together, these data suggest that changes observed in plaques were independent of cholesterol levels.

We next determined key cells present in plaques and interrogated plaques for citrullinated histones (or H3Cit), a marker of extracellular traps that are typically released by neutrophils or macrophages upon activation^24^. Macrophages were determined by the percent of Mac2+ staining and we observed a significant decrease in macrophages in the plaques of young LL mice compared with baseline, suggesting that lipid lowering decreased plaque macrophages (**Figure 1E; Figure S2A**). In contrast to young mice, lipid lowering did not change the percent of plaque macrophages in MA mice. Similar trends were observed with H3Cit, in which there was a decrease in young mice upon lipid lowering compared with baseline (**Figure 1F; Figure S2B**), while lipid lowering had little effect on H3Cit in MA mice. These data demonstrate that middle age limits some effects of lipid lowering on plaque composition/morphology.

An increased fibrous cap is a marker of plaque stability and smooth muscle cells (SMCs) are a major cell type that forms these caps. Therefore, we next examined the levels of alpha SMC actin (αSMA). Young LL mice had increased αSMA relative to baseline, an observation that was not seen in MA mice (**Figure 1G**). In fact, αSMA in MA mice after diet switch was unchanged, and even slightly reduced, compared to the young counterparts (**Figure 1H**). These data suggest that lipid lowering in young mice engages plaque repair mechanisms but that there is a failure to drive these changes in MA mice with lipid lowering therapies.

To further assess inflammation in the plaques, plaque tissue from lipid lowered mice was analyzed by LC-MS/MS for the presence of lipid mediators. Plaques from the aortic arch of MA mice had increased levels of prostanoids (e.g. PGE2 and PGF2A), 12-HHT, and 17-HDHA, compared with young mice after lipid lowering (**Figure S3A-G**). This elevated “prostanoid storm” in MA supports the hypothesis that despite effective cholesterol lowering, MA mice exhibit elevated inflammation.

### Persistent neutrophilia in MA mice despite lipid lowering

Western diet has demonstrated impacts on blood cell production^25,26^. Here, complete blood counts (CBCs) revealed that baseline atherosclerosis was associated with slightly increased red blood cells (RBCs) and reduced platelets in circulation, relative to healthy controls, in both young and MA mice (**Figure S4A-B**). Our analysis also revealed that lipid lowering had no impact on RBCs in either age group, however, MA LL mice, but not young LL mice, exhibited a significant increase in circulating platelets. At baseline, atherosclerosis was associated with increased RBC distribution width (RDW; **Figure S4C**) compared to healthy controls, consistent with clinical data suggesting RDW is associated with progression of atherosclerosis^27^. In both age groups, RDW was significantly reduced upon lipid lowering, consistent with this change being driven by elevated cholesterol. Whereas RBC parameters were regulated primarily by atherosclerosis alone, changes in platelet production upon lipid lowering was more pronounced in MA mice, demonstrating the importance of age as a driver of specific hematological changes.

Atherosclerosis is associated with increased circulating myeloid cells^28,29^, therefore we next investigated the impact of age, atherosclerosis, and lipid lowering on circulating white blood cells (WBCs). Total WBCs were not different in young atherosclerotic mice at baseline or after lipid lowering (**Figure 2A**). However, in MA mice, lipid lowering increased total WBCs relative to baseline. Notably, WBCs were elevated in otherwise healthy MA mice, compared with young healthy controls, but baseline atherosclerosis significantly increased WBCs, suggesting that MA alone did not account for the elevated numbers of WBCs. Monocytes have been found to be critical in both atherosclerosis progression and regression^30,31^. Circulating monocytes were elevated in MA baseline atherosclerotic mice compared with young baseline mice, though this increase was likely driven by age as it was also apparent in healthy age-matched controls (**Figure 2B**). Monocytes were also increased in young baseline atherosclerotic mice relative to age-matched healthy controls, and lipid lowering had no significant impact on circulating monocytes in either young or MA mice. However, lipid lowering induced a significant increase in circulating lymphocytes in both young and MA mice (**Figure 2C**). Together, these data suggest that changes to circulating white blood cells are differentially regulated by both age and diet switch in the context of atherosclerosis.

**Figure 2.**
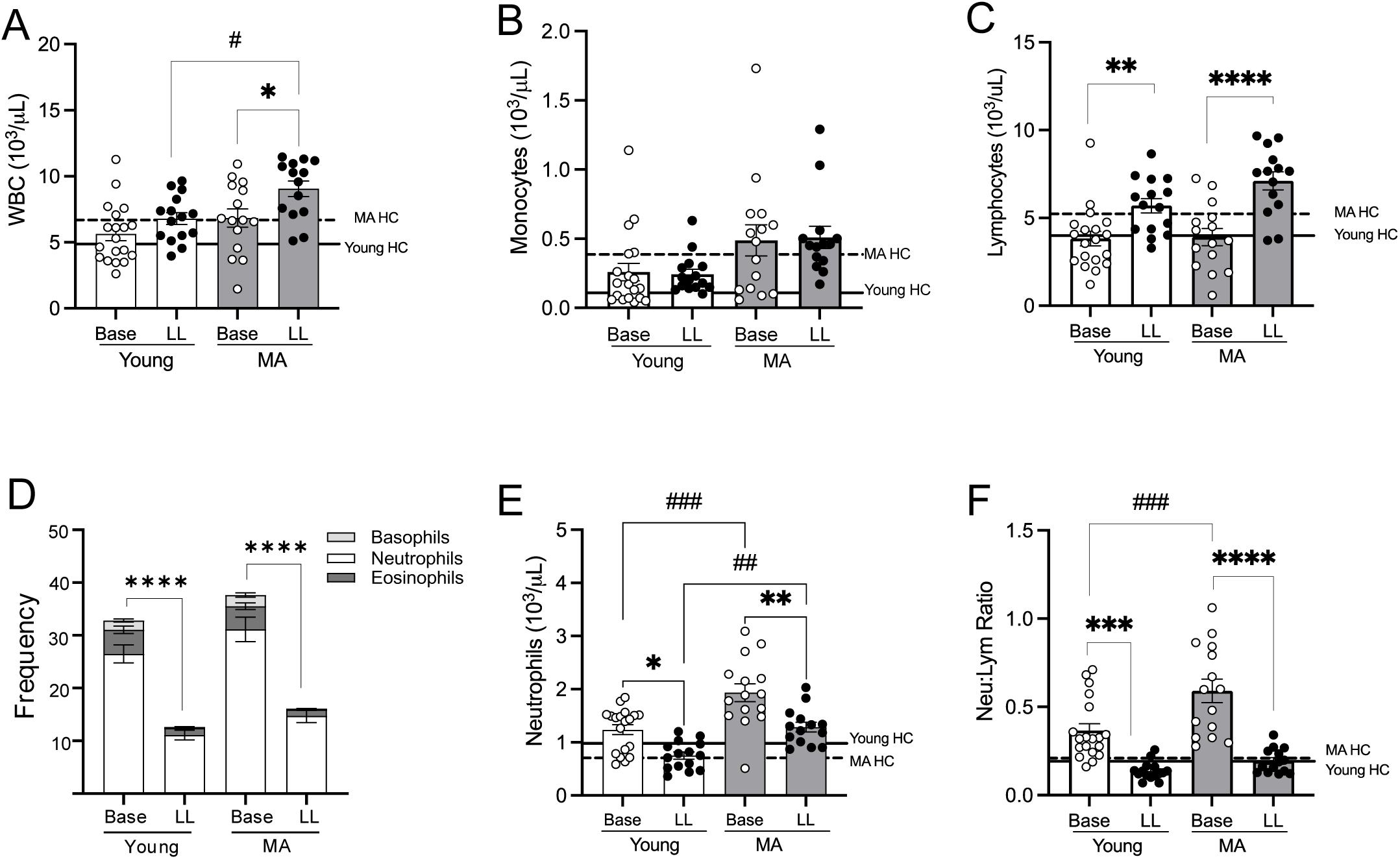
Middle-aged mice exhibit increased circulating myeloid cells despite lipid-lowering. Peripheral blood from mice described in Figure 1 was collected for complete blood counts. Total white blood cells (WBC) (**A**), monocytes (**B**), and lymphocytes (**C**) in young and MA baseline (Base) and lipid lowered (LL) mice is shown. Solid lines reflect mean WBC counts in young healthy controls (Young HC, n=5) and dashed lines depict mean in MA healthy controls (MA HC, n=5). (**D**) Frequencies of granulocytes (neutrophils, eosinophils, and basophils) are shown for young and MA baseline and LL mice. Total neutrophils (**E**) and the neutrophil-to-lymphocyte ratio (**F**) are shown. Data representative of at least 3 independent experiments; n=10-21 mice per group. Results are mean ± SEM. For all analyses, each dot represents an individual mouse and comparisons were performed using two-way ANOVA with Tukey’s multiple comparison. **p<0.01, ***p<0.001, ****p<0.0001 effect of lipid-lowering, #p<0.05 ##p<0.01, ###p<0.001 effect of MA.

Neutrophils are emerging as key cellular players in atherosclerosis^32–34^ and so we next examined circulating granulocytes, which were significantly elevated during atherosclerosis and reduced upon lipid lowering in both age groups (**Figure 2D**). Neutrophils are the most abundant granulocyte in circulation, and we observed significantly more neutrophils in MA baseline mice compared with young baseline (**Figure 2E**). While lipid lowering was sufficient to decrease circulating neutrophils in both young and MA mice, MA LL mice maintained significantly higher circulating neutrophils compared with young LL mice even with lipid-lowering . The neutrophil-to-lymphocyte ratio (NLR) is a clinical marker of inflammation in human atherosclerosis^35^, and MA baseline atherosclerotic mice had a significantly higher ratio than young mice at baseline, though LL diet switch in both young and MA mice was sufficient to reduce the NLR (**Figure 2F**). It is of note that while MA mice showed a reduction in NLR after lipid lowering, this was associated with a significant increase in both circulating lymphocytes and neutrophils after lipid lowering compared to young mice. These results suggest that more advanced disease in MA, even with lipid lowering , are associated with increased WBCs and neutrophils, supporting that aging of the hematopoietic system is a key contributor to atheroprogression.

### Lipid lowering has distinct age-specific impacts on the bone marrow

Because we observed that middle age and lipid lowering induced changes in circulating immune cells, we next examined the bone marrow, the site of blood cell production. Compared to healthy age-matched controls, baseline atherosclerosis resulted in a striking loss of bone marrow cellularity in both young and aged mice (**Figure S5A**), and lipid lowering increased bone marrow cellularity. The lineage-negative, Sca-1^+^ cKit^+^ (LSK) fraction contains stem and progenitor cells and LSK frequencies were significantly increased in MA baseline relative to young baseline , while lipid lowering decreased the LSK pool, in both young and MA (**Figure 3A-B**). As aging alone is associated with an expanded pool of hematopoietic stem cells (HSCs; Lin^-^, Sca1^+^, cKit^+^,CD135^-^, CD150^+^), we next compared phenotypic HSCs and multipotent progenitors in baseline and LL mice^36^, relative to healthy age-matched controls. Frequencies of HSCs were increased in baseline atherosclerosis, relative to healthy, and further increased upon lipid lowering (**Figure 3C**). Multipotent progenitors (MPP) have broad lineage potential similar to HSCs but are more proliferative at homeostasis and support daily blood production. Atherosclerosis led to a significant increase in MPPs in MA relative to young and LL resulted in an expanded pool of MPPs (**Figure 3D**). Numbers of HSCs were increased in MA baseline mice, relative to young baseline atherosclerotic mice, demonstrating that both age and lipid lowering influences the HSC pool (**Figure 3E**). Whereas the entire pool of progenitors was reduced upon lipid lowering in MA mice, the proportion of primitive HSCs and MPPs was expanded, relative to more lineage-committed MPPs.

**Figure 3.**
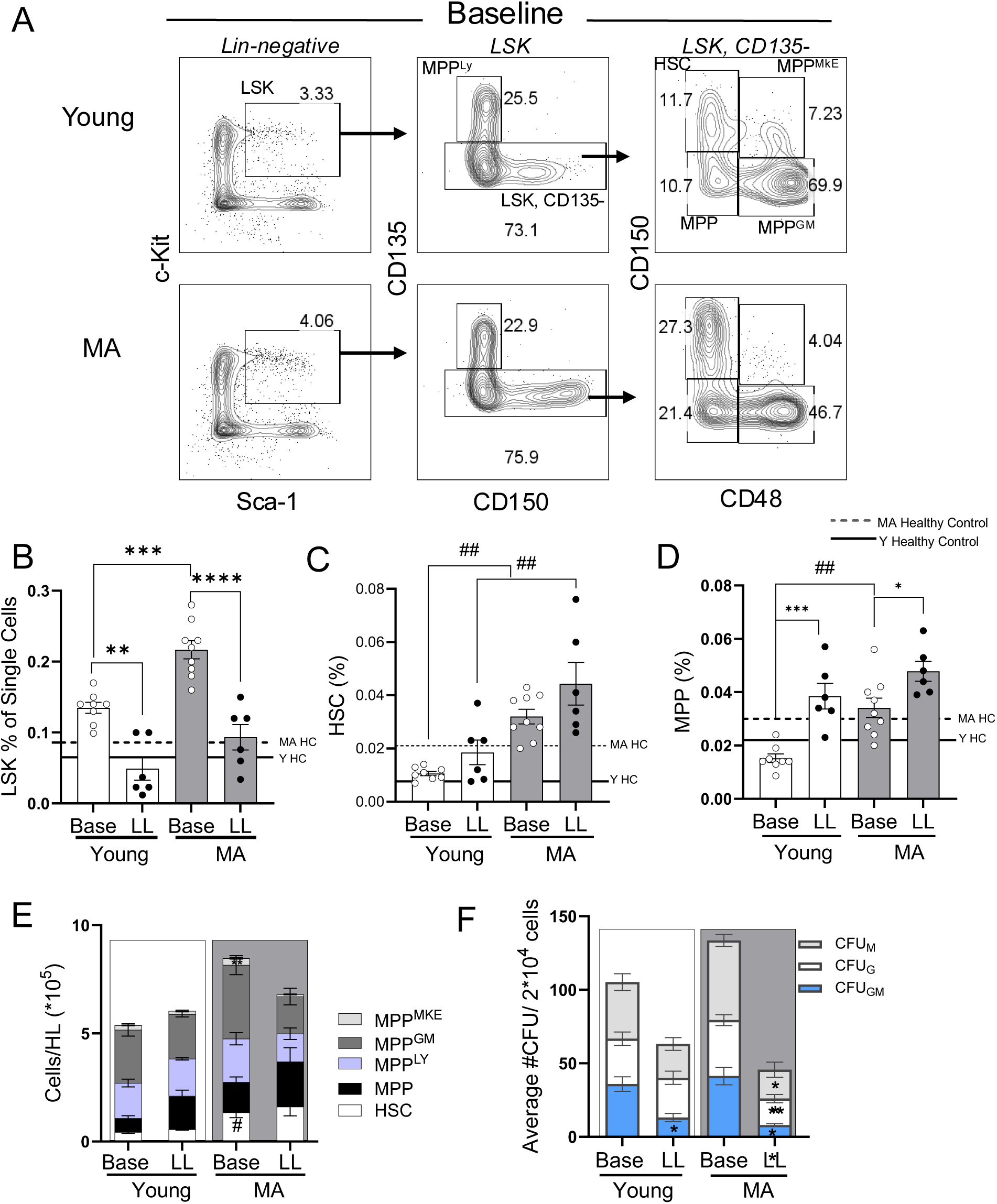
The hematopoietic stem cell pool is impacted by both lipid-lowering and middle age. Bone marrow was collected from the hind limbs of mice treated as in Fig. 1. and single cell suspensions were stained for analysis via flow cytometry to identify hematopoietic stem cells (HSCs) and multipotent progenitors (MPPs). (**A**) The gating schematic used to identify cells is shown for young and middle-aged (MA) mice at baseline. Cells staining positive for lineage markers (CD3, CD11b, Gr-1, Ter119, and B220) were excluded and Lin-negative cells (left column) were analyzed for expression of c-Kit and Sca-1. Lin^-^Sca1^+^c-Kit^+^ (LSK) cells were analyzed for expression of CD135 and CD150 to identify CD135^+^ lymphoid-biased MPPs (MPP^Ly^). The CD135-negative fraction was further analyzed for expression of CD150 and CD48 to identify HSCs, MPPs, MPP^MkE^, and MPP^GM^ (as shown). The numbers adjacent to the gated population reflect the percent among the parent population. (**B**) The absolute frequencies among total bone marrow cells is shown for the LSK population in young and MA baseline (Base) and lipid lowered (LL) mice. (**C-D**) Absolute frequencies of HSCs and MPPs are shown. Data are from 2 independent experiments, n=6-9/group. (**E**) Numbers of each population are shown in young and MA mice at baseline and after lipid lowering. (**F**) Bone marrow cells were plated for colony forming unit (CFU) assays. The numbers of colonies is shown for CFU-G (granulocyte), CFU-M (macrophage, and CFU-GM (granulocyte/ macrophage). Results are mean ± SEM. For all analyses, each dot represents an individual mouse and comparisons were performed using two-way ANOVA with Tukey’s multiple comparison. *p<0.05, **p<0.01, ***p<0.001, ****p<0.0001 effect of lipid-lowering, ##p<0.01, effect of MA.

To evaluate functional differences in the bone marrow, particularly with respect to myeloid cell production, we performed colony forming unit (CFU) assays in MethoCult media. Overall, we identified very few primitive multipotent granulocyte, erythroid, macrophage, megakaryocyte colonies (CFU-GEMM), though more committed granulocyte-macrophage colonies were readily observed. Colonies were slightly increased in MA baseline atherosclerotic mice, compared to young baseline, however lipid lowering reduced colony formation in both young and MA mice (**Figure 3F**). In young, lipid lowering resulted in a significant decrease in primitive CFU-GM but CFU-M and CFU-G colonies were similar between baseline and LL mice. However, in the middle-aged cohort, lipid lowering resulted in a significant decrease both committed CFU-M and CFU-G as well as primitive CFU-GM colonies, suggesting impaired function, relative to young mice. Therefore, despite an expansion of phenotypic HSCs and MPPs in LL MA mice, functional progenitor activity in the bone marrow was markedly reduced in MA upon lipid lowering. As impaired bone marrow function is often associated with extramedullary hematopoiesis, we next evaluated the frequency and function of HSPCs in the spleen. First, we observed that atherosclerotic MA mice having undergone lipid lowering had a greater frequency of splenic HSCs compared with young counterparts (**Figure S5 B, C**). Furthermore, we observed increased CFU-M in the spleens of MA LL mice. These results are consistent with reduced regeneration and repair in the bone marrow during aging and reveal a deficit in bone marrow recovery upon lipid lowering.

### Transcriptional programs driven by lipid lowering are different in young and MA mice

To better understand the global impact of lipid lowering on the bone marrow in young and MA mice, we performed bulk RNAseq on young and MA atherosclerosis mice following lipid lowering treatment. Compared to age-matched healthy controls, MA LL and young LL mice had few shared upregulated genes (**Figure 4A**, red left). Among the shared upregulated genes, *Timd4*, T-cell membrane protein 4, was upregulated in the bone marrow of both young and MA mice following diet switch. There was significant overlap in downregulated genes driven by lipid lowering treatment in young and MA mice which we assessed using Over-Representation Analysis (ORA)^37^ (**Figure 4A**, blue right). Lipid lowering treatment resulted in decreased expression of genes related to matrix remodeling and extracellular structure organization in both young and MA mice, suggesting that these pathways are primarily regulated by diet and not age (**Figure 4B**). Upon lipid lowering, genes involved in VEGF responses and calmodulin binding were downregulated specifically in young mice, but not MA mice, which suggests an improved vascular tone in the bone marrow of young mice with diet switch. In the MA mice, specific genes related to oxidoreductase activity were downregulated upon lipid lowering. Therefore, lipid lowering treatment has distinct transcriptional effects in the bone marrow that depend upon age. We next focused solely on mice having undergone lipid lowering diet switch to directly address the unique signatures in the bone marrow of young and MA mice in this translational context. MA mice exhibited a significant downregulation of 27 genes such as *Gpx2* which is a critical antioxidant, and upregulation of 77 genes, including *Lipin3* and *Pla2g2d* compared with Y controls (**Figure 4C**). To gain mechanistic insight, we applied DoRothEA transcription factor activity analysis, which revealed enhanced activity of Zfp263, Arid3a, and Pou6f2 in MA mice, whereas regulators such as Pax5, Lyl1, and Prdm14 were suppressed (**Figure 4D**). Together, these data suggest that there are distinct transcriptional patterns in the bone marrow between Y an MA LL mice.

**Figure 4.**
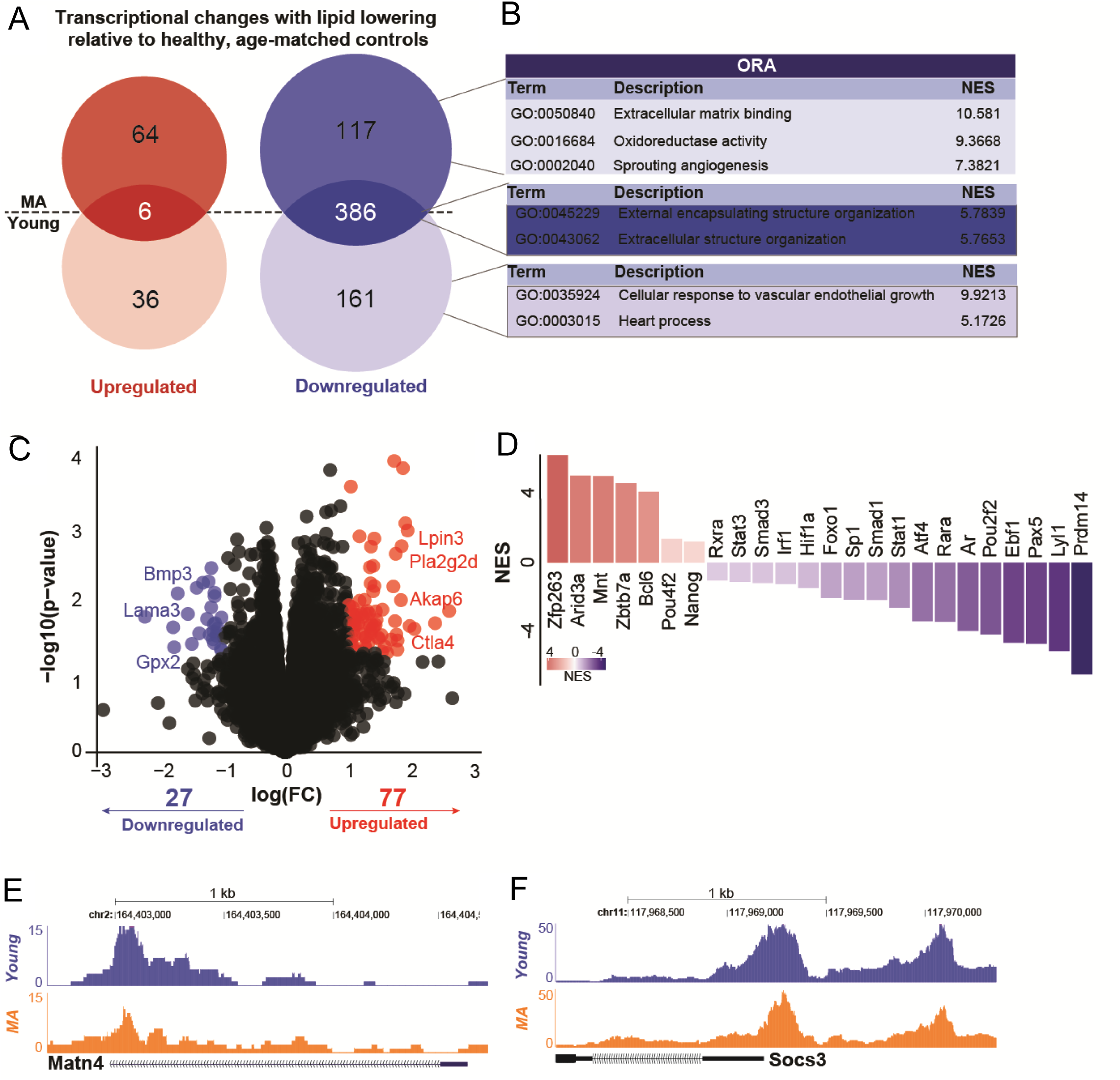
Lipid lowering reveals age-specific transcriptional and chromatin accessibility changes in bone marrow cells. **(A)** Venn diagram of an overlap of significantly upregulated (red) and downregulated (blue) genes in lipid-lowered (LL) young and MA mice relative to age-matched healthy controls. **(B)** Over-representation analysis (ORA, WebGestalt) of downregulated genes in MA mice, highlighting enrichment for extracellular matrix binding, oxidoreductase activity, angiogenesis, and structural organization pathways. **(C)** Volcano plot displaying differentially expressed genes in LL MA mice compared to LL young mice. Selected representative genes are labeled. **(D)** DoRothEA-based transcription factor activity analysis showing transcriptional regulators with significantly increased (red) or decreased (blue) activity in LL MA mice. **(E– F)** ATAC-seq genome browser tracks illustrating age-specific chromatin accessibility.

To test whether age-dependent transcriptional changes were associated with alterations in chromatin accessibility, we performed ATAC-seq on lipid-lowered young and MA mice. This analysis revealed distinct differences in open chromatin regions. For example, *Matn4* showed a pronounced accessibility peak in young mice that was markedly diminished in MA mice (**Figure 4E**). *Socs3* exhibited increased accessibility in young mice compared to MA, consistent with its role as a negative regulator of STAT3 signaling and a key modulator of inflammatory responses (**Figure 4F**). These data suggest that chromatin accessibility in the bone marrow differs between Y and MA mice after lipid lowering. Collectively, lipid lowering is not sufficient to return the MA bone marrow to a state that resembles young bone marrow.

### Neutrophilic inflammation correlates with plaque necrosis in middle-aged atherosclerotic mice despite lipid lowering

Because the bone marrow exhibited several features of elevated inflammation in MA, we next examined mature myeloid cells at this site. We observed that the frequency of bone marrow neutrophils (CD11b^+^, SigF^-^, Ly6C^low^, Ly6G^high^) was significantly increased in young and MA baseline atherosclerosis mice compared with their healthy age-matched controls (**Figure 5A; Figure S6A, B**). We additionally assessed the bone marrow of lipid lowered mice histologically, finding that the total number of Ly6G^+^ cells were higher in lipid lowered MA mice than young mice (**Figure S7A, B**). Similarly to what was observed in the blood, lipid lowering resulted in reduced bone marrow neutrophils. Bone marrow monocytes (CD11b^+^, SigF^-^, Ly6C^high^, Ly6G^low^) were elevated in young baseline atherosclerotic mice and LL significantly reduced this population. In contrast to young mice, frequencies of bone marrow monocytes were unchanged in MA mice when comparing baseline and LL (**Figure 5B**). Baseline atherosclerosis induced an increased proportion of immature CD101^-^ neutrophils, relative to mature CD101^+^ neutrophils^38^, and lipid lowering was able to reduce this slightly (**Figure 5C**). Therefore, lipid lowering treatment exerted a more profound impact on the monocyte lineage of young mice, whereas it exerted more striking changes to the granulocyte compartment of MA mice.

**Figure 5.**
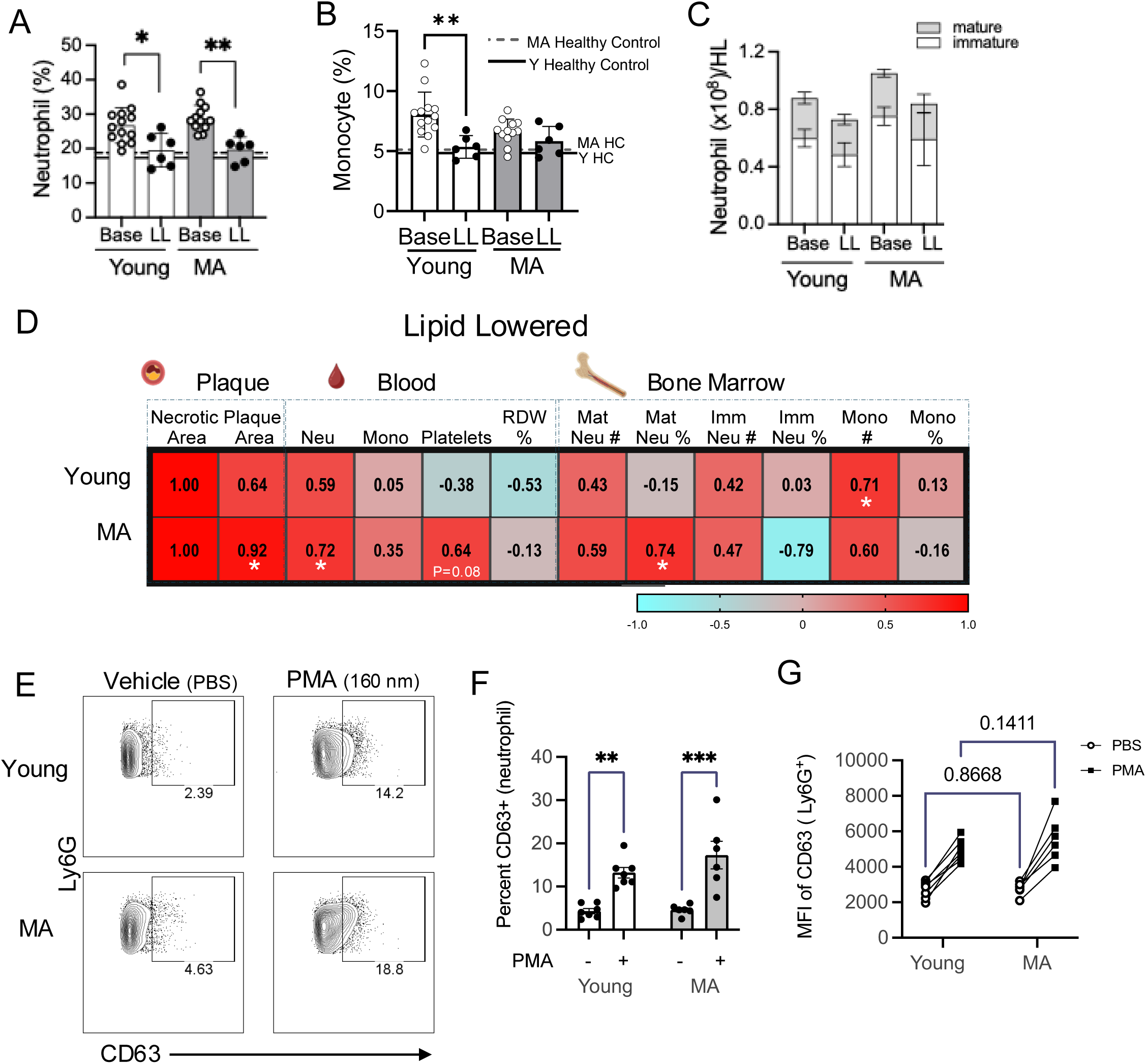
Lipid lowering reduces neutrophilia in both young and MA mice, but neutrophils from MA bone marrow are associated with plaque necrosis. Bone marrow was collected from the hind limbs of mice treated as in Fig. 1. and single cell suspensions were stained for analysis via flow cytometry. (**A**) Neutrophil frequencies (CD11b^+^ SigF^-^ Ly6C^-^ Ly6G^+^) in the bone marrow are shown in young and MA mice at baseline (Base) and upon lipid lowering (LL). (**B**) The frequencies of monocytes (CD11b^+^ SigF^-^ Ly6C^+^ Ly6G^-^) in the bone marrow are shown. Lines on the graphs represent average values of each population in healthy controls. (**C**) The absolute numbers of immature (CD101^-^; white filled) and mature (CD101^+^; gray filled) neutrophils are shown. Data representative of 2 independent experiments; n=6-13 mice per group. Results are mean ± SEM. For all analyses, each dot represents an individual mouse and comparisons were performed using two-way ANOVA with Tukey’s multiple comparison *p<0.05, **p<0.01, effect of lipid-lowering. **(D)** Percent plaque necrosis was correlated with circulating and bone marrow outputs in young and MA LL mice. Pearson’s R for each comparison is shown in the center of the box. Significance is visualized by symbols in the center of the box, *p<0.05. n=6-11/ group. Icons created with Biorender.com (**E**) Representative staining of bone marrow neutrophils (CD11b^+^ SigF^-^ Ly6C^-^ Ly6G^+^) from young and MA LL mice upon stimulation with vehicle or PMA (160nM) showing expression of CD63. The numbers adjacent to the gated population reflect the percent among the parent population. **(F)** The percent of CD63-positive neutrophils is shown in young and MA with vehicle and PMA (100nM). **(G)** The mean fluorescence intensity (MFI) of CD63 on total Ly6G+ cells is shown. Data representative of 2 independent experiments; n=6-7 mice per group. Results are mean ± SEM. For all analyses, each dot represents an individual mouse and comparisons were performed using paired t-test. **p<0.01, ***p<0.0001.

To better understand the complex interactions between age and lipid lowering diet on plaque morphology and hematological changes, we compiled data to perform correlation analyses. We found that the circulating neutrophils were directly and positively correlated with percent plaque necrosis in both young and MA mice, however this was only significant in MA mice when compared using Pearson’s coefficient (**Figure 5D**). Further, the frequency of both immature and mature neutrophils in the bone marrow were significantly correlated with percent plaque necrosis in MA mice only. In contrast to MA mice, plaque necrosis in young LL mice was correlated with the frequency of bone marrow monocytes. Together these findings highlight the link between circulating neutrophil levels, bone marrow composition, and plaque necrosis in MA mice even after effective lipid lowering therapy.

Based on our observation that neutrophils correlated with plaque necrosis in MA LL mice, but not young LL mice, we next addressed potential differences in neutrophil function in young and MA mice. Bone marrow cells were isolated, activated via stimulation with phorbol myristate acetate (PMA) and then analyzed for expression of CD63, a marker of degranulation^39^. Bone marrow neutrophils from both young and MA LL mice had increased CD63 expression in response to PMA stimulation (**Figure 5E**). The percent of CD63+ neutrophils was greater in MA mice (**Figure 5F**). MA mice also had a slightly greater increase in CD63 MFI (mean fluorescence intensity) compared with bone marrow neutrophils from young mice upon lipid lowering (**Figure 5G**). These data suggest that while the frequency of neutrophils may decrease in the bone marrow upon lipid lowering, their activation state is increased in MA relative to young mice, which may account for the direct correlation with increased plaque necrosis in MA.

### Increased neutrophils associated with the brain vasculature of middle-aged mice

Neutrophil adhesion to the cerebral vasculature can contribute to neuroinflammation^40,41^. Given our observation of increased circulating neutrophils in both the contexts of baseline disease as well as following 6 weeks of LL diet, we next examined the effect of LL diet on vascular-associated neutrophils within the hippocampus in young and MA mice. The hippocampus is an integral brain structure involved in learning and memory. Work in humans supports that the hippocampus, and its vasculature in particular, may be particularly sensitive to the effects of chronological age^42^. Furthermore, recent findings have found an association between elevated atherosclerosis pathology in the circle of Willis and hippocampal volume loss^43^. We observed significantly higher lectin (which labels blood vessels) and Ly6G (which labels PMNs) contacts in the hippocampus of MA LL mice compared with young LL mice as measured by immunofluorescence (**Figure 6A, B**), suggesting increased PMN associated with the hippocampal vasculature. We isolated the hippocampus from the brains of LL mice and performed targeted LC-MS/MS analysis and we observed significantly elevated levels of the neutrophil chemoattract leukotriene B_4_ (LTB_4_)^44^ in MA mice (**Figure 6C**). Taken together, our data demonstrate that atherosclerosis may have far-reaching systemic effects, particularly in middle age that may not resolve with lipid lowering.

**Figure 6.**
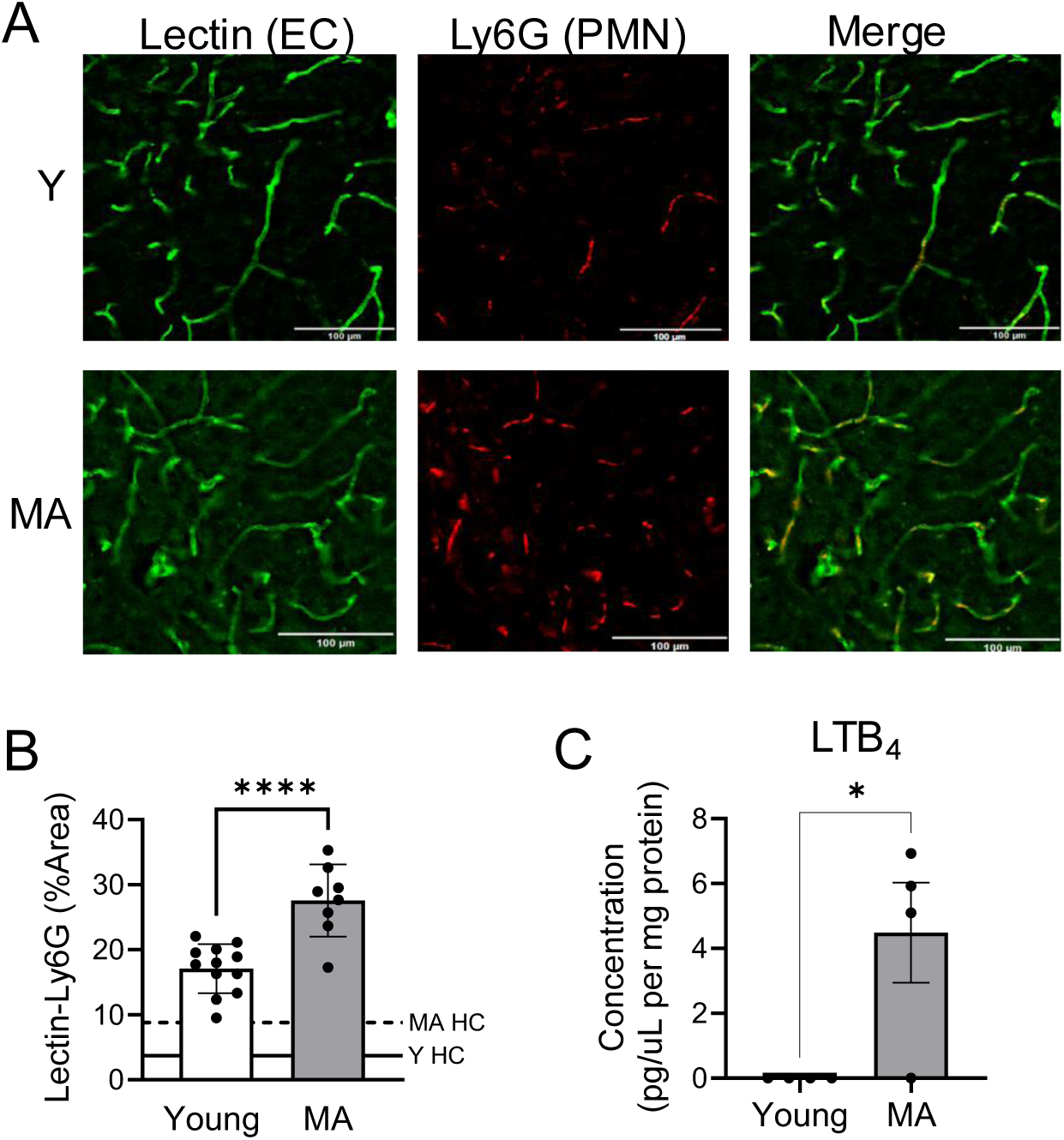
Elevated endothelial-neutrophil contacts in the hippocampus of MA lipid lowered atherosclerotic mice. **(A)** Brains were isolated, sectioned, and immunofluorescence was performed for lectin (green) as a marker of the endothelium and Ly6G (red) as a marker of PMN. **(B)** The percent colocalization of lectin and Ly6G was quantified in the hippocampus using ImageJ. **(C)** The hippocampus was isolated from a subset of randomly selected mice and flash-frozen for LC-MS/MS processing and analysis. Results are mean ± SEM and each dot represents an individual mouse. *p<0.05, ****p<0.0001 Student’s t-test.

## Discussion

We found that MA baseline atherosclerotic mice had similar sized plaques with more necrotic regions, increased PMNs in circulation, and a striking increase in the neutrophil to lymphocyte ratio (NLR), compared with young. Because lipid lowering is the mainstay therapy for atherosclerosis in humans, we asked whether lipid lowering could improve plaque and systemic inflammation equivalently in young and MA mice. Upon lipid lowering we observed a significant increase in necrosis in MA mice, relative to young, that correlated with reduced lesion remodeling, as well as increased circulating WBCs. Increased plaque necrosis in MA upon LL correlated with our observation that MA mice exhibited significantly increased neutrophils and bone marrow neutrophils remained in a more activated state as evidenced by a more robust response to stimulation. In addition, we found that in MA mice neutrophils were also present in the hippocampus, suggesting systemic neutrophilic inflammation driven by atherosclerosis in the MA cohort is not reversed by lipid lowering. These findings support the idea that MA is a critical period of time during which hematopoietic aging coalesces with vascular aging to increase atherosclerosis and hamper the response to lipid-lowering.

Chronological aging has well-characterized impacts on hematopoiesis and HSCs that include an overall myeloid-bias and reduced stem cell function^45,46^. In middle-aged mice, hypercholesterolemia had a more profound ability to prime HSPCs to generate granulocytes. This preferential priming for neutrophils may underlie differences in the progression of atherosclerosis and cellular composition of the plaque in young and middle-aged mice. Indeed, in young mice, hypercholesterolemia elicits inflammatory monocyte programs that drive atherosclerosis^30^ consistent with our observation of an accumulation of macrophages in the plaques of young baseline atherosclerotic mice. Upon lipid lowering, phenotypic HSCs and MPPs were increased, however, consistent with impaired function, we observed reduced CFUs in the bone marrow, a phenomenon that was significantly more pronounced in middle-aged mice. Impaired hematopoiesis in middle age involves not just cell-intrinsic defects^46,47^ but also changes to the stromal microenvironment in the bone marrow^8^. As a result, extramedullary hematopoiesis can occur and splenic hematopoiesis is appreciated to support pathologic myelopoiesis and exacerbate cardiovascular disease^48,49^. Together, our findings suggest that in the context of middle age, reduced regeneration of the bone marrow upon lipid lowering contributes to systemic changes in blood production and splenic function that impact atheroprogression and systemic inflammation. Our findings also support the study of middle-aged mice as way to model translationally relevant aspects of disease and therapeutic approaches.

A key finding herein was that in middle-aged mice, atherosclerosis exhibits a stronger association with neutrophils compared with younger counterparts. While lipid lowering reduces neutrophil abundance, the prolonged circulation of activated neutrophils over several months may foster a maladaptive vascular phenotype distinct from that observed in young mice. Notably, even following lipid lowering, neutrophils persist in a heightened pro-inflammatory state. In agreement, neutrophils are key players in mouse models of atherosclerosis^32,34^. Further corroborating our findings in MA mice, Silvestri-Roig et al also showed a role for neutrophil-associated NET formation in atherosclerotic plaque linked to decreased plaque stability in a modified shear stress model of advanced athersclerosis^50^. We found that despite 6 weeks of lipid lowering, middle-aged mice exhibited increased plaque necrosis and NET formation compared to young mice on the same regiment. In humans, neutrophilia has been recognized as an independent predictor of adverse events in patients with acute coronary syndrome^33,51^.

Additional clinical findings such as the neutrophil-to-lymphocyte ratio (NLR) have consistently been shown to predict cardiovascular risk and all-cause mortality in analyses of data from multiple randomized trials^35^. Analysis of two patient cohorts (n=1570) with established arterial disease identified a panel of 50 proteins linked to neutrophil-driven inflammatory pathways^52^. Collectively, these findings in relation to ours highlight neutrophil-driven inflammation as a lasting contributor to residual risk and support its potential as a biomarker for heightened atherosclerotic risk. Our data also suggests that lipid lowering can reduce the NLR in both Y and MA mice, however, bone marrow neutrophils exhibited a more heightened responsiveness to PMA stimulation in MA mice, suggesting the potential for a more pro-inflammatory neutrophil phenotype. Together, these data support the idea that treating atherosclerosis in middle-age may require additional complementary strategies to limit PMN reactivity.

The above concept becomes especially critical given the new link we found between neutrophils and the hippocampal vasculature in atherosclerosis. Atherosclerosis and dementia are leading causes of mortality and morbidity worldwide^53,54^. Atherosclerosis increases dementia risk 2.5 fold^55^ and is considered a major, independent risk factor for vascular contributions to cognitive impairment and dementia (VCID) ^53^. Neutrophil adhesion to the brain vasculature has been associated with endothelial activation upon brain injury and may contribute to neurovascular inflammation^40,41,56^. Moreover, neutrophils have also been implicated in neuroinflammatory diseases including ischemic stroke and Alzheimer’s disease, where neutrophils were shown to associate with capillary stalls leading to reduced cerebral blood flow^41,57–59^. Blockade of dynamic capillary stall formation with anti-Ly6G antibody showed benefit in an ischemic reperfusion model in mice^58^. Work herein suggests that despite lipid lowering, atherosclerosis in middle age elevates neutrophils associated with the hippocampal vasculature and further work will be required to define the mechanistic relationship between neuroinflammation in the hippocampus, and atherosclerosis. Collectively, these data suggest that additional and complementary therapies that control systemic inflammation, without suppressing immunity, are needed to limit the non-resolving systemic inflammation during atherosclerosis in middle age.

## Supporting information

Gannon Rahtes Supplemental Figures

## Acknowledgements

This work was supported by NIH grant HL170249 (G.F., K.C.M.) Special thanks to Christina Thrasher (Albany Medical College, Albany, NY) for her assistance with tissue processing and optimization.

## Author contributions

O.G. and A.R. performed and analyzed experiments and wrote the manuscript. J.B. and I.S.D.R. and R.B.R performed and analyzed experiments and contributed to writing of the manuscript. J.P., S.K, A.R., G.R.C., A.B. performed and analyzed experiments. K.C.M. and G.F. conceived and designed experiments, analyzed experiments, wrote the manuscript and provided funding.

## Notes

### Competing Interest Statement

The authors have declared no competing interest.

## References

1 Bohula, E. A. et al. Achievement of dual low-density lipoprotein cholesterol and high-sensitivity C-reactive protein targets more frequent with the addition of ezetimibe to simvastatin and associated with better outcomes in IMPROVE-IT. Circulation 132, 1224–1233 (2015). 10.1161/CIRCULATIONAHA.115.018381

2 Pradhan, A. D., Aday, A. W., Rose, L. M. & Ridker, P. M. Residual Inflammatory Risk on Treatment With PCSK9 Inhibition and Statin Therapy. Circulation 138, 141–149 (2018). 10.1161/CIRCULATIONAHA.118.034645

3 Lopez-Melgar, B. et al. Short-Term Progression of Multiterritorial Subclinical Atherosclerosis. J Am Coll Cardiol 75, 1617–1627 (2020). 10.1016/j.jacc.2020.02.026

4 Wong, M. Y. Z., Yap, J., Huang, W., Tan, S. Y. & Yeo, K. K. Impact of Age and Sex on Subclinical Coronary Atherosclerosis in a Healthy Asian Population. JACC Asia 1, 93–102 (2021). 10.1016/j.jacasi.2021.05.002

5 Fredman, G. & Serhan, C. N. Specialized proresolving mediator targets for RvE1 and RvD1 in peripheral blood and mechanisms of resolution. Biochem J 437, 185–197 (2011). BJ20110327 [pii]10.1042/BJ20110327

6 Libby, P. Inflammation during the life cycle of the atherosclerotic plaque. Cardiovasc Res 117, 2525–2536 (2021). 10.1093/cvr/cvab303

7 Rohde, D. et al. Bone marrow endothelial dysfunction promotes myeloid cell expansion in cardiovascular disease. Nat Cardiovasc Res 1, 28–44 (2022). 10.1038/s44161-021-00002-8

8 Young, K. et al. Decline in IGF1 in the bone marrow microenvironment initiates hematopoietic stem cell aging. Cell Stem Cell 28, 1473–1482 e1477 (2021). 10.1016/j.stem.2021.03.017

9 Fitzgerald, H. et al. Resolvin D2-GPR18 Enhances Bone Marrow Function and Limits Steatosis and Hepatic Collagen Accumulation in Aging. Am J Pathol (2023). 10.1016/j.ajpath.2023.08.011

10 Du, W. et al. Age-associated vascular inflammation promotes monocytosis during atherogenesis. Aging Cell 15, 766–777 (2016). 10.1111/acel.12488

11 Boucher, D. M. et al. Age-Related Impairments in Immune Cell Efferocytosis and Autophagy Hinder Atherosclerosis Regression. Arterioscler Thromb Vasc Biol 45, 481–495 (2025). 10.1161/ATVBAHA.124.321662

12 Keeter, W. C., Ma, S., Stahr, N., Moriarty, A. K. & Galkina, E. V. Atherosclerosis and multi-organ-associated pathologies. Semin Immunopathol 44, 363–374 (2022). 10.1007/s00281-022-00914-y

13 Gustavsson, A. M. et al. Midlife Atherosclerosis and Development of Alzheimer or Vascular Dementia. Ann Neurol 87, 52–62 (2020). 10.1002/ana.25645

14 Cortes-Canteli, M. et al. Subclinical Atherosclerosis and Brain Metabolism in Middle-Aged Individuals: The PESA Study. J Am Coll Cardiol 77, 888–898 (2021). 10.1016/j.jacc.2020.12.027

15 Gottesman, R. F. et al. Associations Between Midlife Vascular Risk Factors and 25-Year Incident Dementia in the Atherosclerosis Risk in Communities (ARIC) Cohort. JAMA Neurol 74, 1246–1254 (2017). 10.1001/jamaneurol.2017.1658

16 Liang, J. et al. Associations Between Atherosclerosis and Subsequent Cognitive Decline: A Prospective Cohort Study. J Am Heart Assoc 13, e036696 (2024). 10.1161/JAHA.124.036696

17 Buchanan, H., Hull, C., Cacho Barraza, M., Delibegovic, M. & Platt, B. Apolipoprotein E loss of function: Influence on murine brain markers of physiology and pathology. Aging Brain 2, 100055 (2022). 10.1016/j.nbas.2022.100055

18 Seidel, F. et al. Ldlr-/-.Leiden mice develop neurodegeneration, age-dependent astrogliosis and obesity-induced changes in microglia immunophenotype which are partly reversed by complement component 5 neutralizing antibody. Front Cell Neurosci 17, 1205261 (2023). 10.3389/fncel.2023.1205261

19 Rodrigues, M. S. et al. Microglia contribute to cognitive decline in hypercholesterolemic LDLr(-/-) mice. J Neurochem 168, 1565–1586 (2024). 10.1111/jnc.15952

20 Lipscomb, M. et al. Resolvin D2 limits atherosclerosis progression via myeloid cell-GPR18. FASEB J 38, e23555 (2024). 10.1096/fj.202302336RR

21 Amend, S. R., Valkenburg, K. C. & Pienta, K. J. Murine Hind Limb Long Bone Dissection and Bone Marrow Isolation. J Vis Exp (2016). 10.3791/53936

22 Damascena, H. L., Silveira, W. A. A., Castro, M. S. & Fontes, W. Neutrophil Activated by the Famous and Potent PMA (Phorbol Myristate Acetate). Cells 11 (2022). 10.3390/cells11182889

23 Zhao, Y. et al. Stage-specific remodeling of atherosclerotic lesions upon cholesterol lowering in LDL receptor knockout mice. Am J Pathol 179, 1522–1532 (2011). S0002-9440(11)00519-0 [pii]10.1016/j.ajpath.2011.05.020

24 Mauracher, L. M. et al. Citrullinated histone H3, a biomarker of neutrophil extracellular trap formation, predicts the risk of venous thromboembolism in cancer patients. J Thromb Haemost 16, 508–518 (2018). 10.1111/jth.13951

25 Benson, T. W. et al. A single high-fat meal provokes pathological erythrocyte remodeling and increases myeloperoxidase levels: implications for acute coronary syndrome. Lab Invest 98, 1300–1310 (2018). 10.1038/s41374-018-0038-3

26 Unruh, D. et al. Red Blood Cell Dysfunction Induced by High-Fat Diet: Potential Implications for Obesity-Related Atherosclerosis. Circulation 132, 1898–1908 (2015). 10.1161/CIRCULATIONAHA.115.017313

27 Lappegard, J. et al. Red cell distribution width and carotid atherosclerosis progression. The Tromso Study. Thromb Haemost 113, 649–654 (2015). 10.1160/TH14-07-0606

28 Poller, W. C., Nahrendorf, M. & Swirski, F. K. Hematopoiesis and Cardiovascular Disease. Circ Res 126, 1061–1085 (2020). 10.1161/CIRCRESAHA.120.315895

29 Louloudis, G. et al. Adeno-Associated Virus-Mediated Gain-of-Function mPCSK9 Expression in the Mouse Induces Hypercholesterolemia, Monocytosis, Neutrophilia, and a Hypercoagulative State. Front Cardiovasc Med 8, 718741 (2021). 10.3389/fcvm.2021.718741

30 Swirski, F. K. et al. Ly-6Chi monocytes dominate hypercholesterolemia-associated monocytosis and give rise to macrophages in atheromata. J Clin Invest 117, 195–205 (2007). 10.1172/JCI29950

31 Rahman, K. et al. Inflammatory Ly6Chi monocytes and their conversion to M2 macrophages drive atherosclerosis regression. J Clin Invest 127, 2904–2915 (2017). 10.1172/JCI75005

32 Lavillegrand, J. R. et al. Alternating high-fat diet enhances atherosclerosis by neutrophil reprogramming. Nature 634, 447–456 (2024). 10.1038/s41586-024-07693-6

33 Soehnlein, O. & Doring, Y. Beyond association: high neutrophil counts are a causal risk factor for atherosclerotic cardiovascular disease. Eur Heart J 44, 4965–4967 (2023). 10.1093/eurheartj/ehad711

34 Drechsler, M., Megens, R. T., van Zandvoort, M., Weber, C. & Soehnlein, O. Hyperlipidemia-triggered neutrophilia promotes early atherosclerosis. Circulation 122, 1837–1845 (2010). 10.1161/CIRCULATIONAHA.110.961714

35 Adamstein, N. H. et al. The neutrophil-lymphocyte ratio and incident atherosclerotic events: analyses from five contemporary randomized trials. Eur Heart J 42, 896–903 (2021). 10.1093/eurheartj/ehaa1034

36 Challen, G. A., Pietras, E. M., Wallscheid, N. C. & Signer, R. A. J. Simplified murine multipotent progenitor isolation scheme: Establishing a consensus approach for multipotent progenitor identification. Exp Hematol 104, 55–63 (2021). 10.1016/j.exphem.2021.09.007

37 Elizarraras, J. M. et al. WebGestalt 2024: faster gene set analysis and new support for metabolomics and multi-omics. Nucleic Acids Res 52, W415–W421 (2024). 10.1093/nar/gkae456

38 Gullotta, G. S. et al. Age-induced alterations of granulopoiesis generate atypical neutrophils that aggravate stroke pathology. Nat Immunol 24, 925–940 (2023). 10.1038/s41590-023-01505-1

39 Simard, J. C., Girard, D. & Tessier, P. A. Induction of neutrophil degranulation by S100A9 via a MAPK-dependent mechanism. J Leukoc Biol 87, 905–914 (2010). 10.1189/jlb.1009676

40 Smyth, L. C. D. et al. Neutrophil-vascular interactions drive myeloperoxidase accumulation in the brain in Alzheimer’s disease. Acta Neuropathol Commun 10, 38 (2022). 10.1186/s40478-022-01347-2

41 Cruz Hernandez, J. C., et al. Neutrophil adhesion in brain capillaries reduces cortical blood flow and impairs memory function in Alzheimer’s disease mouse models. Nat Neurosci 22, 413–420 (2019). 10.1038/s41593-018-0329-4

42 Montagne, A. et al. Blood-brain barrier breakdown in the aging human hippocampus. Neuron 85, 296–302 (2015). 10.1016/j.neuron.2014.12.032

43 Kapasi, A. et al. Atherosclerosis and Hippocampal Volumes in Older Adults: The Role of Age and Blood Pressure. J Am Heart Assoc 13, e031551 (2024). 10.1161/JAHA.123.031551

44 Afonso, P. V. et al. LTB4 is a signal-relay molecule during neutrophil chemotaxis. Dev Cell 22, 1079–1091 (2012). 10.1016/j.devcel.2012.02.003

45 Beerman, I. et al. Functionally distinct hematopoietic stem cells modulate hematopoietic lineage potential during aging by a mechanism of clonal expansion. Proc Natl Acad Sci U S A 107, 5465–5470 (2010). 10.1073/pnas.1000834107

46 Chambers, S. M. et al. Aging hematopoietic stem cells decline in function and exhibit epigenetic dysregulation. PLoS Biol 5, e201 (2007). 10.1371/journal.pbio.0050201

47 Kuribayashi, W. et al. Limited rejuvenation of aged hematopoietic stem cells in young bone marrow niche. J Exp Med 218 (2021). 10.1084/jem.20192283

48 Robbins, C. S. et al. Extramedullary hematopoiesis generates Ly-6C(high) monocytes that infiltrate atherosclerotic lesions. Circulation 125, 364–374 (2012). CIRCULATIONAHA.111.061986[pii]10.1161/CIRCULATIONAHA.111.061986

49 Dutta, P. et al. E-Selectin Inhibition Mitigates Splenic HSC Activation and Myelopoiesis in Hypercholesterolemic Mice With Myocardial Infarction. Arterioscler Thromb Vasc Biol 36, 1802–1808 (2016). 10.1161/ATVBAHA.116.307519

50 Silvestre-Roig, C. et al. Externalized histone H4 orchestrates chronic inflammation by inducing lytic cell death. Nature 569, 236–240 (2019). 10.1038/s41586-019-1167-6

51 Guasti, L. et al. Neutrophils and clinical outcomes in patients with acute coronary syndromes and/or cardiac revascularisation. A systematic review on more than 34,000 subjects. Thromb Haemost 106, 591–599 (2011). 10.1160/TH11-02-0096

52 Nurmohamed, N. S. et al. Targeted proteomics improves cardiovascular risk prediction in secondary prevention. Eur Heart J 43, 1569–1577 (2022). 10.1093/eurheartj/ehac055

53 Collaborators, G. B. D. D. F. Estimation of the global prevalence of dementia in 2019 and forecasted prevalence in 2050: an analysis for the Global Burden of Disease Study 2019. Lancet Public Health 7, e105–e125 (2022). 10.1016/S2468-2667(21)00249-8

54 Martin, S. S. et al. 2025 Heart Disease and Stroke Statistics: A Report of US and Global Data From the American Heart Association. Circulation 151, e41–e660 (2025). 10.1161/CIR.0000000000001303

55 Wendell, C. R. et al. Carotid atherosclerosis and prospective risk of dementia. Stroke 43, 3319–3324 (2012). 10.1161/STROKEAHA.112.672527

56 Herz, J. et al. Role of Neutrophils in Exacerbation of Brain Injury After Focal Cerebral Ischemia in Hyperlipidemic Mice. Stroke 46, 2916–2925 (2015). 10.1161/STROKEAHA.115.010620

57 Santos-Lima, B., Pietronigro, E. C., Terrabuio, E., Zenaro, E. & Constantin, G. The role of neutrophils in the dysfunction of central nervous system barriers. Front Aging Neurosci 14, 965169 (2022). 10.3389/fnagi.2022.965169

58 Erdener, S. E. et al. Dynamic capillary stalls in reperfused ischemic penumbra contribute to injury: A hyperacute role for neutrophils in persistent traffic jams. J Cereb Blood Flow Metab 41, 236–252 (2021). 10.1177/0271678X20914179

59 Crumpler, R., Roman, R. J. & Fan, F. Capillary Stalling: A Mechanism of Decreased Cerebral Blood Flow in AD/ADRD. J Exp Neurol 2, 149–153 (2021). 10.33696/neurol.2.048

