## Supplementary material for "Lipid lowering alone fails to limit atherosclerosis progression and neutrophilic inflammation in middle-aged mice": Gannon Rahtes Supplemental Figures

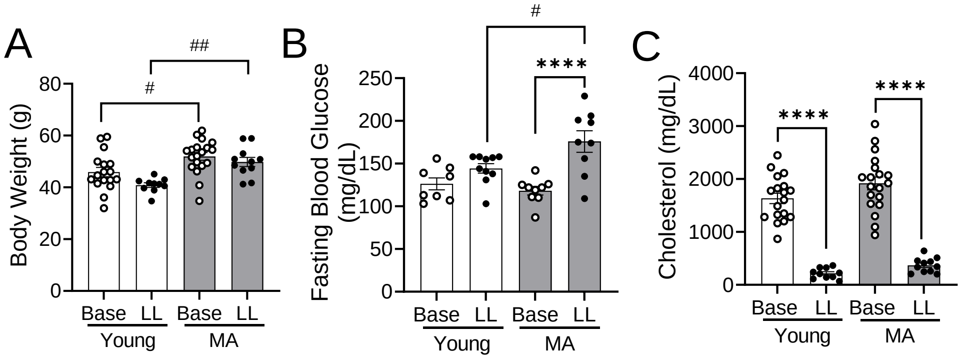


**Supplemental Figure S1**. **Age drives differences in weight and fasting glucose, but not cholesterol.** Experiments were performed as in Fig 1. **(A)** Body weight (in grams) is shown. **(B)** One week prior to sacrifice, mice were fasted for 6 hours and blood glucose was measured from blood samples taken via tail vein using a Trumetrix glucometer. **(C)** Total plasma cholesterol at the time of sacrifice was measured as described in the methods. Data are representative of at least 3 independent experiments; n=10-19 mice per group. Results are mean ± SEM. For all analyses, each dot represents an individual mouse and comparisons were performed using two-way ANOVA with Tukey's multiple comparison. ****$p<0.0001$ effect of lipid-lowering**,** #$p<0.05$ ##$p<0.01$ effect of MA.

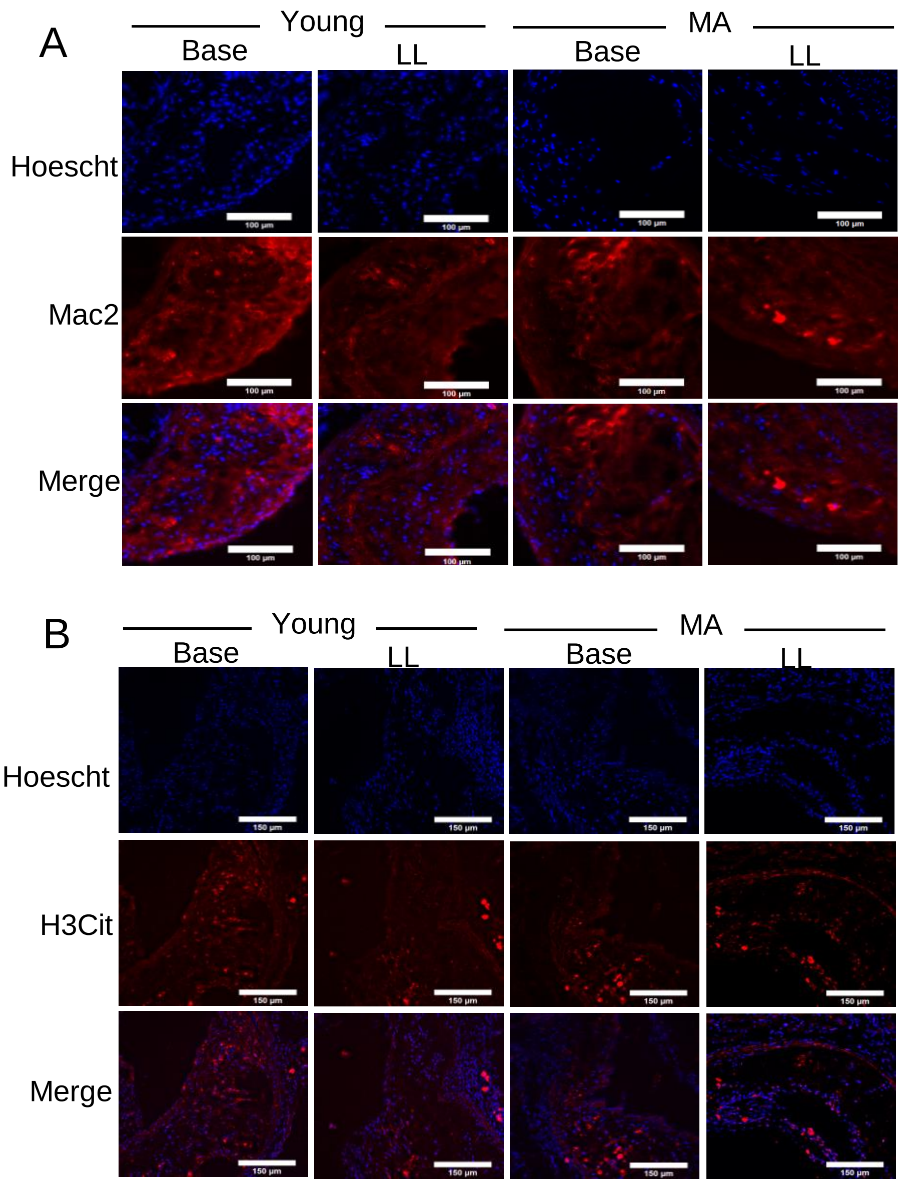


**Supplemental Figure S2**. Experiments were performed as in Fig 1. (**A**) Representative images of Mac2 (red) expression in plaques collected from either young (Y) or middle-aged (MA) mice at either baseline (Base = 20 weeks post western diet) or after lipid lowering (LL = 20 weeks western diet + 6 weeks chow diet). (**B)** Representative images of citrullinated histone (red) expression in plaques collected from Y and MA mice at either baseline or after lipid lowering. For all images, individual nuclei were stained with Hoescht (blue) and images were acquired on a Thunder microscope.

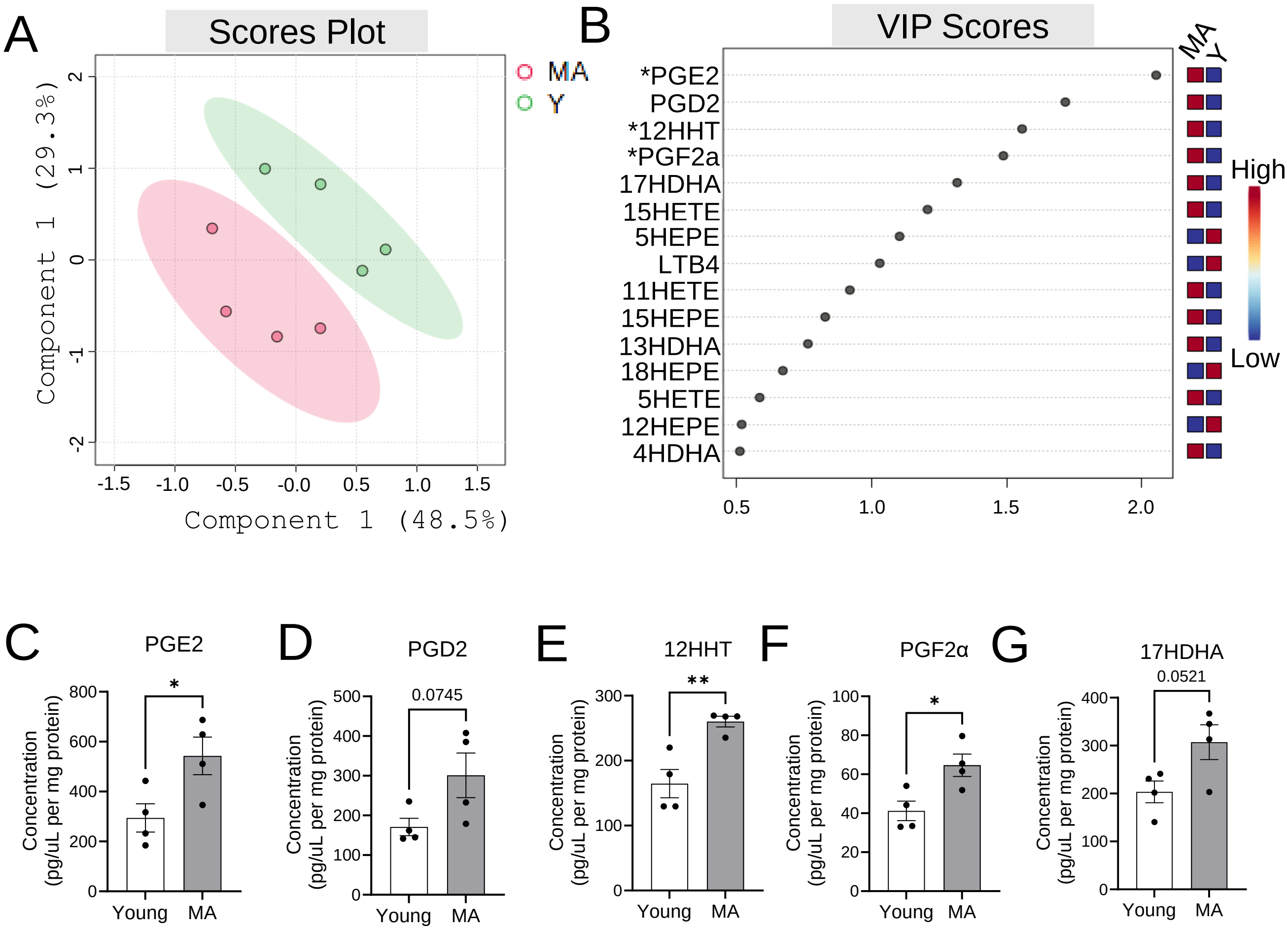


**Supplemental Figure S3. Plaques of middle-aged mice show distinct lipid mediator profiles compared to young mice following lipid lowering therapy.** Experiments were performed as in Fig 1. **A)** PLS-DA of plaque-associated aortic arch lipid mediator profiles in young (Y, green) and middle-aged (MA, red) mice post lipid lowering. **B)** Variable importance projection (VIP) scores. Lipid mediators at the right of 1 are important contributors to the PLS-DA plot, while those at or below 1 are less important. Those that are increased relative to young mice are indicated with red squares, while those that are decreased are indicated with blue squares (on the right). Component 1: R2=0.8, Q2=0.2;Component 2: R2=0.8 ,Q2=-0.1. **C-G)** Each detected lipid mediator was subjected to an independent t-test analysis between Y and MA. The top 5 lipid mediators on the VIP score plot are shown with their significance (as assessed by t-test comparison). Results are representative of one experiment, where each dot indicates an individual mouse.*p<0.05, **p<0.01. PLS-DA and VIP score analysis was performed using Metaboanalyst using log-transformed normalization of the raw concentration values. t-test was used for comparison of the raw concentration value. All data is representative of n=8 (4=Y, 4=MA) from one independent experiment.

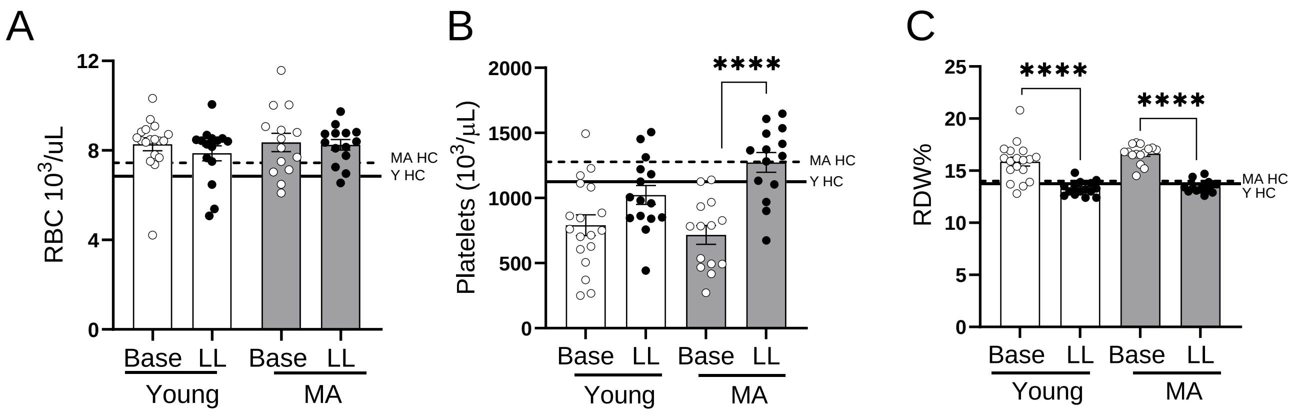


**Supplemental Figure S4. Hematological changes in young and MA baseline and lipid lowered mice.** Peripheral blood from experimental mice described in Figure 1 was collected for complete blood counts. (**A**) Red blood cells (RBC), (**B**) platelets, and (**C**) RBC distribution width (RDW%) are shown in young and MA baseline (Base) and lipid lowered (LL) mice is shown. Solid lines reflect mean values of each parameter in young healthy controls (Young HC, n=5) and dashed lines depict mean values in MA healthy controls (MA HC, n=5). Data are representative of at least 3 independent experiments; n=14-18 mice per group. Results are mean ± SEM. For all analyses, each dot represents an individual mouse and comparisons were performed using two-way ANOVA with Tukey's multiple comparison. ****p<0.0001 effect of lipid-lowering.

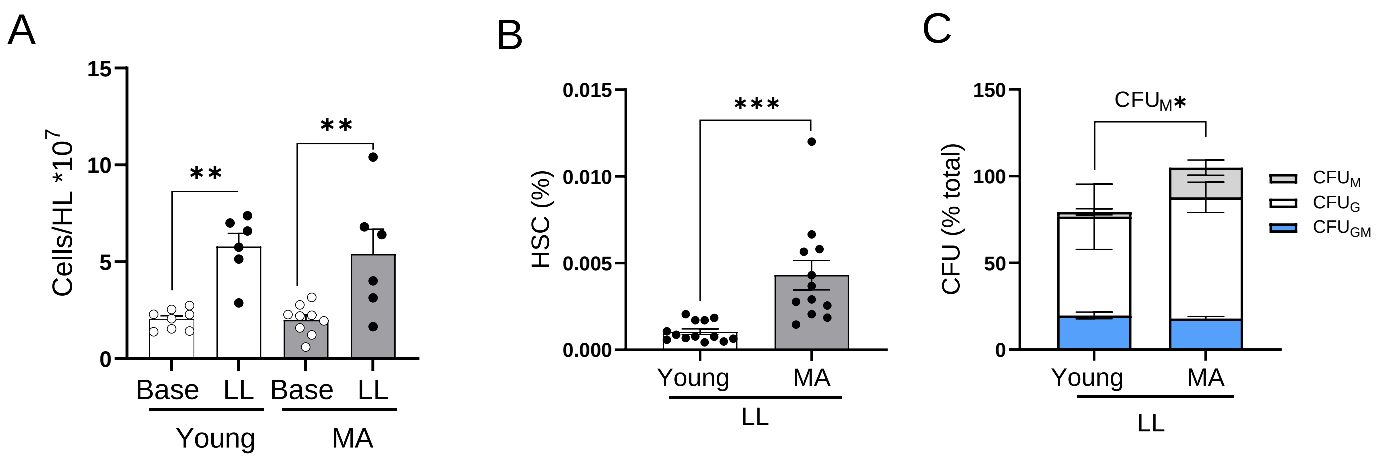


**Supplemental Figure S5. Hematopoietic progenitors from the bone marrow and the spleen are functionally different in lipid lowered MA mice.** Bone marrow was collected from the hind limbs of mice as in Fig. 1. Cells were stained **(A)** Hind limb (HL) bone marrow cellularity was counted, excluding dead cells. Data are from 2 independent experiments, n=6-9/group. Results are mean ± SEM. Analyses were performed using two-way ANOVA with Tukey's multiple comparison. **p<0.01 effect of lipid-lowering. **(B)** Spleens were harvested from young and MA lipid lowered mice and processed. Single cell suspensions were stained for analysis via flow cytometry to identify hematopoietic stem cells (HSCs) using the gating strategy outlined in figure 3A. Cells staining positive for lineage markers (CD3, CD11b, Gr-1, Ter119, and B220) were excluded and Lin-negative cells were analyzed for expression of c-Kit and Sca-1. $Lin^-Sca1^+c\text{-}Kit^+$ (LSK) cells were analyzed for expression of CD135 and CD150. The $CD135^-$ fraction was analyzed for expression of CD150 and CD48 to identify HSCs. Data are from 1 experiment, n=12-14/group. Results are mean ± SEM. Analyses were performed using T-test. ***p<0.001. For all analyses, each dot represents an individual mouse. **(C)** Spleen cells plated for colony forming unit assay and are displayed as a percentage of total colonies. Colonies were identified between day 6 and 7. Data are from 1 experiment, n=6/group. Analyses were performed using T-test. *p<0.05.

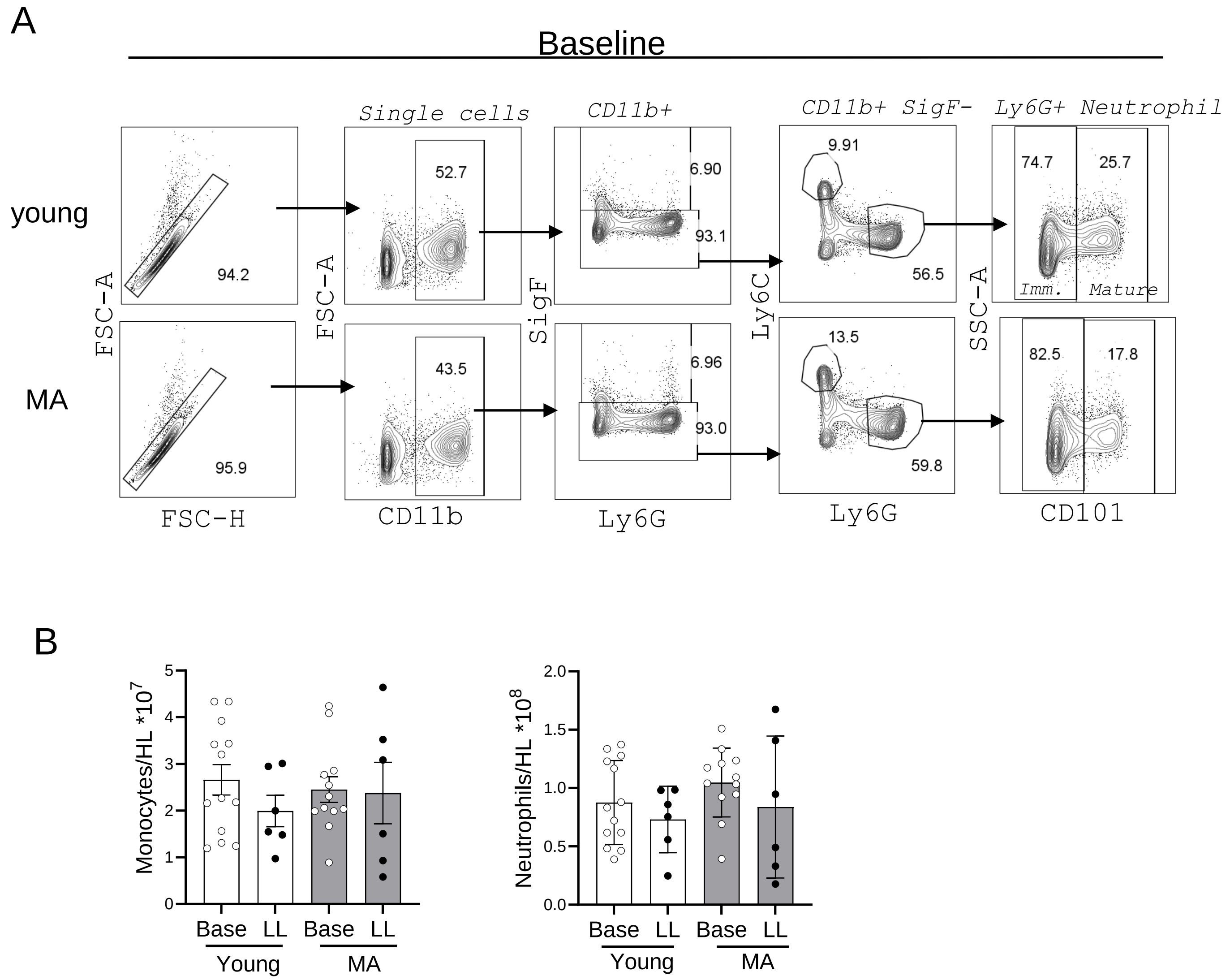


**Supplemental Figure S6. Bone marrow neutrophils and monocytes.** Bone marrow was collected from the hind limbs (HLs) of mice treated as in Fig. 1. and single cell suspensions were stained for analysis via flow cytometry to identify bone marrow monocytes and neutrophils. (**A**) The gating schematic used to identify cells is shown for young (top) and middle-aged (MA; bottom) mice at baseline. Single cells positive CD11b$^+$ and lacking SigF expression were analyzed for Ly6G and Ly6C expression. (**B**) Numbers of each population are shown in young and MA mice at baseline and after lipid lowering. Neutrophil and monocyte numbers are shown**.** Data are from 2 independent experiments, n=6-13/group. Data are representative of 2 independent experiments; n=6-13 mice per group. Results are mean ± SEM. For all analyses, each dot represents an individual mouse and comparisons were performed using two-way ANOVA with Tukey's multiple comparison *p<0.05, **p<0.01, effect of lipid-lowering.

A

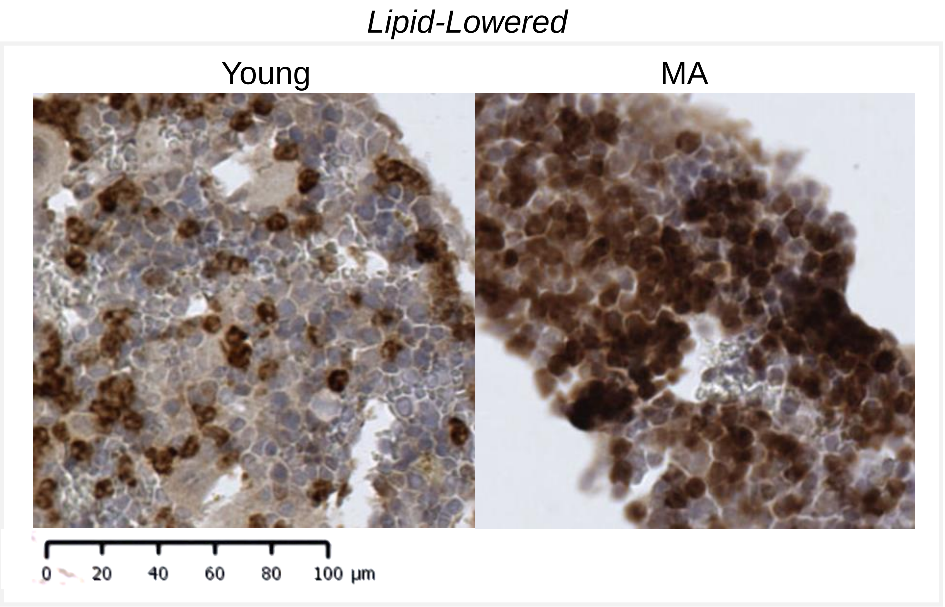


B

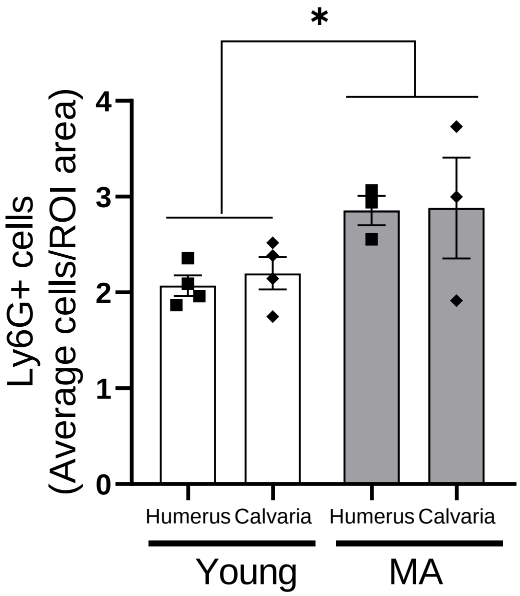


**Supplemental Figure S7 (A-B). Ly6G+ cells in different bone marrow locations.** The humerus and calvaria were dissected and processed for paraffin embedding and sectioning. Average Ly6G$^+$ cells were quantified by a blinded observer. Data representative of 1 experiment; n=3-4 mice per group. Results are mean ± SEM. For analyses, each dot represents an individual mouse and comparisons were performed using paired two-way ANOVA with Tukey's multiple comparison *p<0.05 effect of MA.
